# TIP60 acetylates H2AZ and regulates doxorubicin-induced DNA damage sensitivity through *RAD51* transcription

**DOI:** 10.1101/2020.06.10.145193

**Authors:** Kwok Kin Lee, Yanzhou Zhang, Roberto Tirado- Magallanes, Deepa Rajagopalan, Shreshtha Sailesh Bhatia, Larry Ng, Ng Desi, Cheng Yong Tham, Wen Shiun Teo, Michal Marek Hoppe, Anand D. Jeyasekharan, Yvonne Tay, Wee Joo Chng, Daniel G. Tenen, Touati Benoukraf, Sudhakar Jha

## Abstract

TIP60, a lysine acetyltransferase and H2AZ, a histone H2A variant are involved in transcription and DNA repair. Recent studies suggest that H2AZ acetylation is dependent on TIP60. Here, we show that TIP60 acetylates both isoforms of H2AZ *in vitro* and in cells. Utilizing ChIP-seq and RNA-seq to identify the genes regulated by TIP60-dependent acetylation of H2AZ, we find that TIP60-dependent acetylation of H2AZ correlates with the expression of genes involved in DNA damage repair, amongst several other pathways. In line with this, TIP60-depleted cells exhibit increased sensitivity to the DNA damage-inducing drug doxorubicin. Restoring the expression level of *RAD51*, one of the genes involved in the DNA damage repair pathway, partially rescues the doxorubicin sensitivity due to TIP60 depletion. Overall, our study uncovers a role for TIP60 in regulating doxorubicin-induced DNA damage sensitivity in a manner dependent on *RAD51* transcription.

## Introduction

The nucleosome core particle (NCP) is the basic unit of eukaryotic chromatin consisting of approximately 147-200 bp of DNA wrapped around a histone octamer (Harp et al, 2000; Kornberg, 1974; Luger et al, 1997). Within the histone octamer resides two subunits each of canonical histones H2A, H2B, H3, and H4. Each histone comprises two components namely, the globular core domain which constitutes the histone bulk, and the unstructured tail domain which frequently undergoes posttranslational modifications (PTMs) (Mersfelder & Parthun, 2006). Histone tail PTMs are involved in chromatin remodeling and play important roles in transcription regulation (Mersfelder & Parthun, 2006). Among the PTMs on histone tails, acetylation on lysine residues of histones and its role in transcription regulation is well established (Clayton et al, 2006; Jenuwein & Allis, 2001; Kuo & Allis, 1998; Sterner & Berger, 2000; Struhl, 1998; Workman & Kingston, 1998). Histone acetylation neutralizes lysine residue charges, disrupting histone-DNA interactions, resulting in an open chromatin structure which aids the accessibility of transcription components to the underlying DNA for transcription activation (Clayton et al, 2006; Jenuwein & Allis, 2001; Kuo & Allis, 1998; Sterner & Berger, 2000; Struhl, 1998; Workman & Kingston, 1998).

Acetylation marks are deposited by a class of enzymes termed histone acetyltransferases (HAT) (Sterner & Berger, 2000). An example of a HAT is Tat-interactive protein 60kDa (TIP60), an acetyltransferase within the MYST (<u>M</u>oz, <u>Y</u>bf2/Sas3, <u>S</u>as2, and <u>T</u>IP60) family (Avvakumov & Cote, 2007; Sapountzi et al, 2006). TIP60 has been shown to acetylate histones H3, H4, and H2A (Kimura & Horikoshi, 1998; Yamamoto & Horikoshi, 1997) and through its acetyltransferase activity, TIP60 regulates multiple cellular activities such as transcription, DNA repair, and cell cycle (Brady et al, 1999; Frank et al, 2003; Hejna et al, 2008; Ikura et al, 2015; Ikura et al, 2000; Jacquet et al, 2016; Jeong et al, 2011; Jha et al, 2010; Kusch et al, 2004; Li et al, 2019; Miyamoto et al, 2008; Murr et al, 2006; Sapountzi et al, 2006; Squatrito et al, 2006; Su et al, 2017; Sun et al, 2005; Sun et al, 2010; Tang et al, 2013; Taubert et al, 2004; Van Den Broeck et al, 2012). As a transcription co-activator, TIP60 is recruited by various transcription factors or nuclear receptors to the promoter of cellular genes for the acetylation of histones (Brady et al, 1999; Frank et al, 2003; Jeong et al, 2011; Taubert et al, 2004). In the DNA damage repair context, TIP60 is recruited to sites of DNA damage for chromatin remodeling and histone acetylation, which regulates the loading of DNA repair factors to DNA double-strand break sites (Ikura et al, 2015; Jacquet et al, 2016; Kusch et al, 2004; Li et al, 2019; Murr et al, 2006; Squatrito et al, 2006; Su et al, 2017; Sun et al, 2010; Tang et al, 2013). TIP60 also aids the DNA repair process through the acetylation of DNA damage repair protein ATM, which increases ATM’s catalytic activity leading to activation of downstream DNA damage repair signaling (Sun et al, 2005). Furthermore, loss of TIP60 increases cellular sensitivity to various DNA damage agents, indicating the importance of TIP60 in DNA damage repair (Hejna et al, 2008; Ikura et al, 2000; Miyamoto et al, 2008; Su et al, 2017; Sun et al, 2005; Tang et al, 2013). Apart from direct regulation at DNA damage sites, a few studies found that several DNA repair genes are downregulated upon TIP60 depletion, which could link TIP60’s transcription regulation function to DNA damage sensitivity (Miyamoto et al, 2008; Su et al, 2017; Van Den Broeck et al, 2012).

H2AZ is a histone H2A variant with approximately 60% similarity to canonical histone H2A (Suto et al, 2000). It is highly conserved across various organisms and is essential for the viability of *Drosophila, Tetrahymena*, and mouse (Faast et al, 2001; Santisteban et al, 2000). In humans, there are 2 non-allelic isoforms of H2AZ, namely H2AZ.1 and H2AZ.2, expressed from the *H2AFZ* and *H2AFV* loci respectively (Dryhurst et al, 2009; Horikoshi et al, 2013; Matsuda et al, 2010). Although the two isoforms differ by only three amino acids, their underlying nucleotide sequences are distinct. H2AZ is involved in diverse cellular roles ranging from transcription, DNA damage repair, telomeric functions and chromosome segregation (Babiarz et al, 2006; Dalvai et al, 2013a; Dalvai et al, 2013b; Domaschenz et al, 2017; Farris et al, 2005; Gevry et al, 2007; Giaimo et al, 2018; Hu et al, 2013; John et al, 2008; Pradhan et al, 2016; Rangasamy et al, 2004; Santisteban et al, 2000; Segala et al, 2016; Sutcliffe et al, 2009; Valdes-Mora et al, 2012; Vardabasso et al, 2015; Weber et al, 2010; Xu et al, 2012; Zlatanova & Thakar, 2008).

In the transcription context, H2AZ is enriched at promoters, enhancers and regulatory regions of cellular genes, and is implicated in both activation and repression of genes (Barski et al, 2007; Brunelle et al, 2015; Dalvai et al, 2013a; Dalvai et al, 2013b; Domaschenz et al, 2017; Farris et al, 2005; Gevry et al, 2007; Giaimo et al, 2018; Halley et al, 2010; Hu et al, 2013; John et al, 2008; Law & Cheung, 2015; Pradhan et al, 2016; Santisteban et al, 2000; Segala et al, 2016; Sutcliffe et al, 2009; Valdes-Mora et al, 2017; Valdes-Mora et al, 2012; Vardabasso et al, 2015; Zlatanova & Thakar, 2008). This differential regulation is likely dependent on the type and combination of PTMs on H2AZ (Giaimo et al, 2018; Sevilla & Binda, 2014; Subramanian et al, 2015). Among the PTMs, acetylation on H2AZ has consistently been shown to promote gene transcription. H2AZ is acetylated on lysines 4, 7, 11, 13 and 15, and its hyperacetylated form has been shown to associate with active genes in chicken cells by affecting nucleosome stability and promoting an open conformation (Bonenfant et al, 2006; Bruce et al, 2005; Dryhurst et al, 2009; Ishibashi et al, 2009; Law & Cheung, 2015; Millar et al, 2006; Sevilla & Binda, 2014). Likewise in yeast, the acetylated H2AZ homolog HTZ1 has been shown to associate with the promoter of active genes genome-wide to influence gene activity (Halley et al, 2010; Keogh et al, 2006; Mehta et al, 2010; Millar et al, 2006). In humans, the correlation between acetylated H2AZ and active transcription has been reported, and the wide-spread deregulation of acetylated H2AZ occupancy might play a role in the context of cancer (Dalvai et al, 2013a; Giaimo et al, 2018; Hu et al, 2013; Valdes-Mora et al, 2017; Valdes-Mora et al, 2012).

H2AZ has also been reported to play a direct role in DNA damage repair through chromatin remodeling around the DNA damage site (Xu et al, 2012). Upon DNA damage, H2AZ is rapidly incorporated into DNA damage sites to disrupt nucleosome stability and to promote chromatin acetylation and ubiquitination for the loading of DNA repair factors (Gursoy-Yuzugullu et al, 2015; Xu et al, 2012). Correspondingly, loss of H2AZ results in sensitivity to irradiation-induced DNA damage, indicating the importance of H2AZ in DNA damage repair (Xu et al, 2012). However, whether H2AZ plays a transcriptional role in DNA damage repair is unknown.

Homologous recombination (HR) is the cell’s error-free DNA repair mechanism to fix DNA double-strand breaks through the use of a homologous template for repair (Sullivan & Bernstein, 2018). Loss of HR leads to the utilization of alternative repair pathways, such as non-homologous end joining (NHEJ) or alt-NHEJ, to repair DNA (Tang et al, 2013; Xu et al, 2012). These repair mechanisms are usually error-prone which could result in genomic instability such as chromosome aberrations or loss, leading to apoptosis or carcinogenesis (Li & Heyer, 2008; Tang et al, 2013; Xu et al, 2012). Depletion of H2AZ or TIP60 results in a decrease in HR repair and subsequent increase in alternative DNA repair pathways, leading to cellular sensitivity to DNA damage (Jacquet et al, 2016; Li et al, 2019; Su et al, 2017; Tang et al, 2013; Xu et al, 2012).

In yeast, the H2AZ homolog HTZ1 is deposited and acetylated by the TIP60 homologous complexes – SWR1 and NuA4 – respectively (Altaf et al, 2010; Babiarz et al, 2006; Keogh et al, 2006; Krogan et al, 2004; Krogan et al, 2003; Mehta et al, 2010; Millar et al, 2006; Mizuguchi et al, 2004). Similarly, in *Drosophila*, the dTIP60 complex acetylates H2AV, which is homologous to both the human H2AZ and H2AX, and aids in the exchange of unmodified H2AV for phosphorylated H2AV into chromatin (Kusch et al, 2004). In humans, several lines of evidence suggest that TIP60 is the HAT which acetylates H2AZ (Choi et al, 2009; Dalvai et al, 2013a; Giaimo et al, 2018; Sevilla & Binda, 2014). In this study, we explore genes regulated by TIP60-dependent H2AZ acetylation using a genome-wide approach. We first confirm that TIP60 acetylates H2AZ both *in vitro* and in the cellular context. Subsequently, utilizing chromatin immunoprecipitation followed by sequencing (ChIP-seq) and RNA sequencing (RNA-seq), we identified several genes involved in homologous DNA repair, amongst other pathways, which could be regulated through this mechanism. *RAD51*, one of the genes involved in the DNA repair pathway, was validated and shown to be important in regulating TIP60-dependent cellular resistance to DNA damage.

## Results

### TIP60 acetylates H2AZ *in vitro* and in the cellular context

TIP60 is known to acetylate canonical histones H2A at lysine 5 and H4 at lysines 5, 8, 12 and 16 (Kimura & Horikoshi, 1998; Yamamoto & Horikoshi, 1997). Consistent with previous reports that H2AZ is evolutionarily conserved (Santisteban et al, 2000; van Daal et al, 1990), alignment of human H2AZ.1 or H2AZ.2, with H2AZ of other species shows high homology (Fig. 1A), suggesting an important functional role played by H2AZ across species. H2AZ is acetylated at lysines 4, 7, 11, 13 and 15 (Bonenfant et al, 2006; Bruce et al, 2005; Dryhurst et al, 2009; Ishibashi et al, 2009; Law & Cheung, 2015; Millar et al, 2006; Sevilla & Binda, 2014). As a preliminary study, we align the known H2AZ acetylation sites with TIP60-dependent H4 and H2A acetylation sites (Fig. 1A). We note that H2A lysine 5 aligns with lysine 7 of both H2AZ.1 and H2AZ.2 (Fig. 1A). In addition, known TIP60 acetylation sites on H4 lysines 5, 8, 12 and 16 align perfectly with H2AZ acetylation sites on lysines 4, 7, 11 and 15 for both human H2AZ isoforms respectively (Fig. 1A). The lysine residues of H2AZ.1 and H2AZ.2 are also perfectly aligned with each other and with H2AZ of other species (Fig. 1A). These observations suggest that TIP60 might be able to acetylate both human H2AZ isoforms at these lysines.

**Fig. 1.**
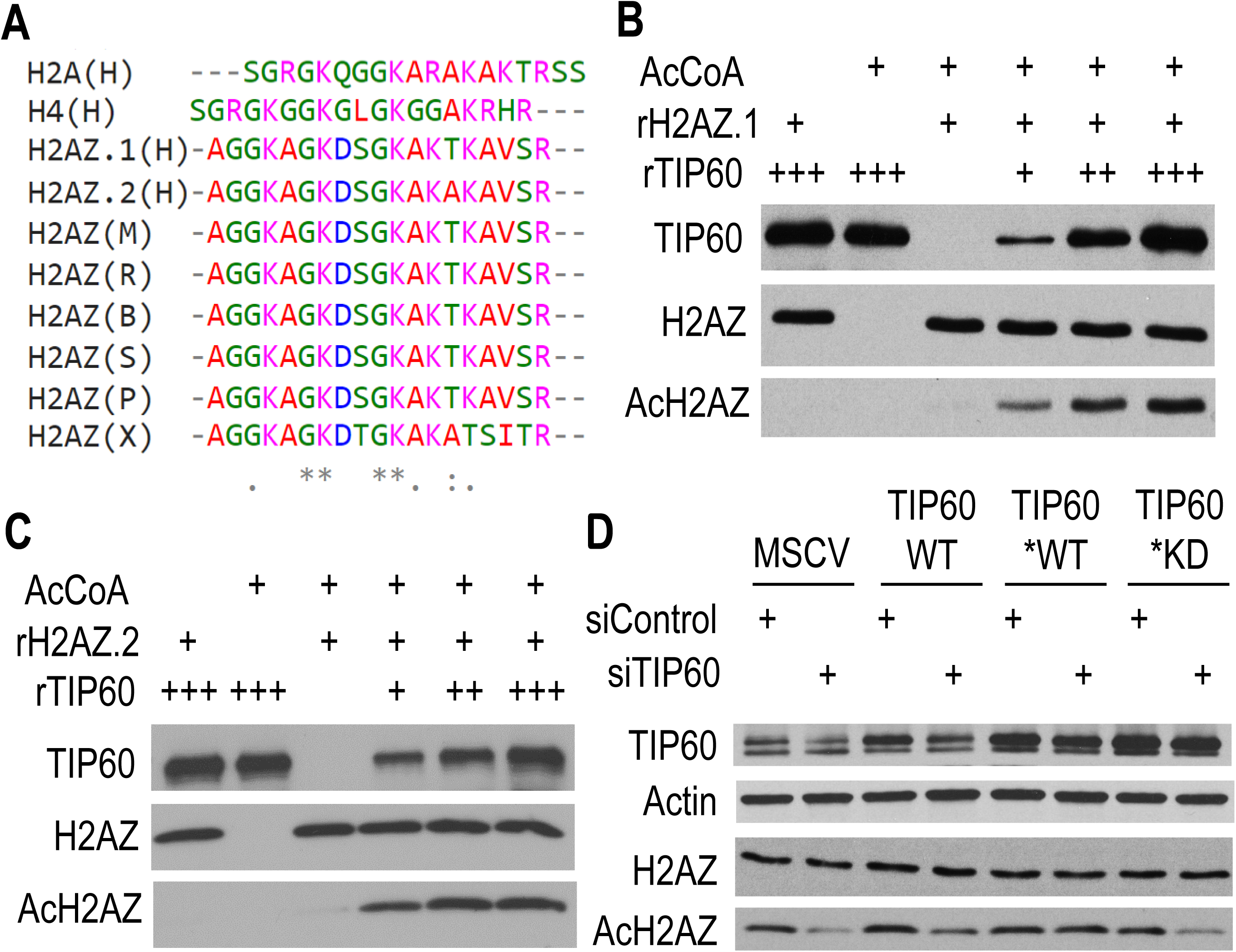
TIP60 acetylates H2AZ *in vitro* and in the cellular context. (**A**) Alignment of H2AZ sequences from multiple species to human histone H4 and H2A sequence. H, human; M, mouse; R, rat; B, bovine; S, sheep; P, pongo; X, *Xenopus*. (**B**) *In vitro* HAT assay using recombinant TIP60 and H2AZ.1 showing TIP60 directly acetylates H2AZ.1. 2000 ng of rH2AZ.1 and 0 ng, 500 ng, 1000 ng and 2000 ng of TIP60 were used. r, recombinant; AcCoA, acetyl-coenzyme A. (N = 2). (**C**) *In vitro* HAT assay using recombinant TIP60 and H2AZ.2 showing TIP60 directly acetylates H2AZ.2. 2000 ng of rH2AZ.2 and 0 ng, 500 ng, 1000 ng and 2000 ng of TIP60 were used. r, recombinant; AcCoA, acetyl-coenzyme A. (N = 3). (**D**) Acetylated H2AZ in MCF10A cells decreases upon depletion of TIP60 and is rescued by TIP60 wild-type but not by the catalytic inactive form. *, siTIP60 resistant; WT, wild-type; KD, catalytic inactivate. Actin serves as a loading control. (N = 3).

To show that H2AZ is a direct substrate of TIP60, an *in vitro* histone acetyltransferase activity (HAT) assay is performed using recombinant TIP60 (rTIP60) and H2AZ.1 (rH2AZ.1) or H2AZ.2 (rH2AZ.2). To detect acetylated H2AZ, we use an AcH2AZ antibody which recognizes acetylated lysines 4, 7 and 11 of H2AZ (AcH2AZ). The same AcH2AZ and H2AZ antibodies were used by a previous study to show a positive correlation between AcH2AZ on promoters and active gene expression (Valdes-Mora et al, 2012). In the presence of acetyl-CoA, TIP60 acetylates both rH2AZ.1 and rH2AZ.2, as seen by the presence of an AcH2AZ band when all three components (rH2AZ.1/2, TIP60, and AcCoA) are present (Fig. 1B and 1C). This also indicates that the AcH2AZ and H2AZ antibodies used can detect both isoforms of acetylated and total H2AZ respectively. Acetylation of H2AZ.1 and H2AZ.2 are specific to TIP60 as observed from a dose-dependent increase in AcH2AZ upon increasing amounts of TIP60 (Fig. 1B and 1C). This data is consistent with previous reports in yeast and *Drosophila* where purified TIP60 complex acetylates either free H2AZ, H2AZ in octamer, or chromatin H2AZ (Altaf et al, 2010; Kusch et al, 2004). Furthermore, the H2AZ complex containing TIP60 purified from the human cell line has been shown to acetylate H2AZ in octamer and in nucleosome *in vitro* (Choi et al, 2009), indicating that TIP60 can acetylate both free H2AZ, as shown in our study, as well as on chromatin.

To validate this observation in the cellular context, we perform siRNA depletion of TIP60 in MCF10A cells stably expressing vector control (MSCV). AcH2AZ level decreases upon the transfection of siTIP60 (Fig. 1D). The specificity of this enzyme-substrate combination is further confirmed by a rescue experiment where we transfected siTIP60 into different MCF10A stable cell lines (Fig. 1D). AcH2AZ level in MCF10A stably expressing siTIP60-resistant wild-type TIP60 (TIP60*WT) is rescued to control level compared to MCF10A MSCV, while a partial rescue is observed for non-resistant wild-type TIP60 (TIP60WT) (Fig. 1D). Furthermore, expressing siTIP60-resistant catalytic inactive TIP60 (TIP60*KD) failed to rescue the AcH2AZ level upon TIP60 depletion (Fig. 1D), indicating that H2AZ acetylation is dependent on TIP60’s catalytic activity. These results suggest that the acetylation of H2AZ is specific to TIP60 and dependent on TIP60’s catalytic activity.

### TIP60 regulates acetylation and occupancy of H2AZ at transcription start site (TSS)

Although TIP60 has been shown to regulate AcH2AZ on the transcription start site (TSS) of some genes (Dalvai et al, 2013a; Giaimo et al, 2018), no genome-wide study has been done to identify all genes regulated by this mechanism. Here, we perform genome-wide ChIP-seq and RNA-seq in MCF10A cells treated with either siControl or siTIP60.

ChIP-seq analysis reveals that the majority of AcH2AZ and H2AZ peaks (47.9% and 42.4% respectively) affected by TIP60 depletion are localized on gene promoters (TSS +/- 2kb) (Fig. S1A and S1B). We also note that both H2AZ and AcH2AZ at transcription start site (TSS) are enriched in a bimodal distribution (Fig. 2A and 2B), similar to what others have reported (Barski et al, 2007; Raisner et al, 2005; Valdes-Mora et al, 2012). On gene promoters, depletion of TIP60 results in major abrogation of both H2AZ and acetylated H2AZ occupancy (Fig. 2A and 2B). Similar occupancy changes are observed on Nucleolin (*NUC*) but not *ACHR* promoter (known positive and negative TIP60 occupied sites, respectively) (Frank et al, 2003; Jha et al, 2010), further supporting changes in H2AZ acetylation and occupancy on TIP60 occupied TSS (Fig. S2A and S2B). These results suggest that TIP60 promotes H2AZ and AcH2AZ occupancy on gene promoters.

**Fig. 2.**
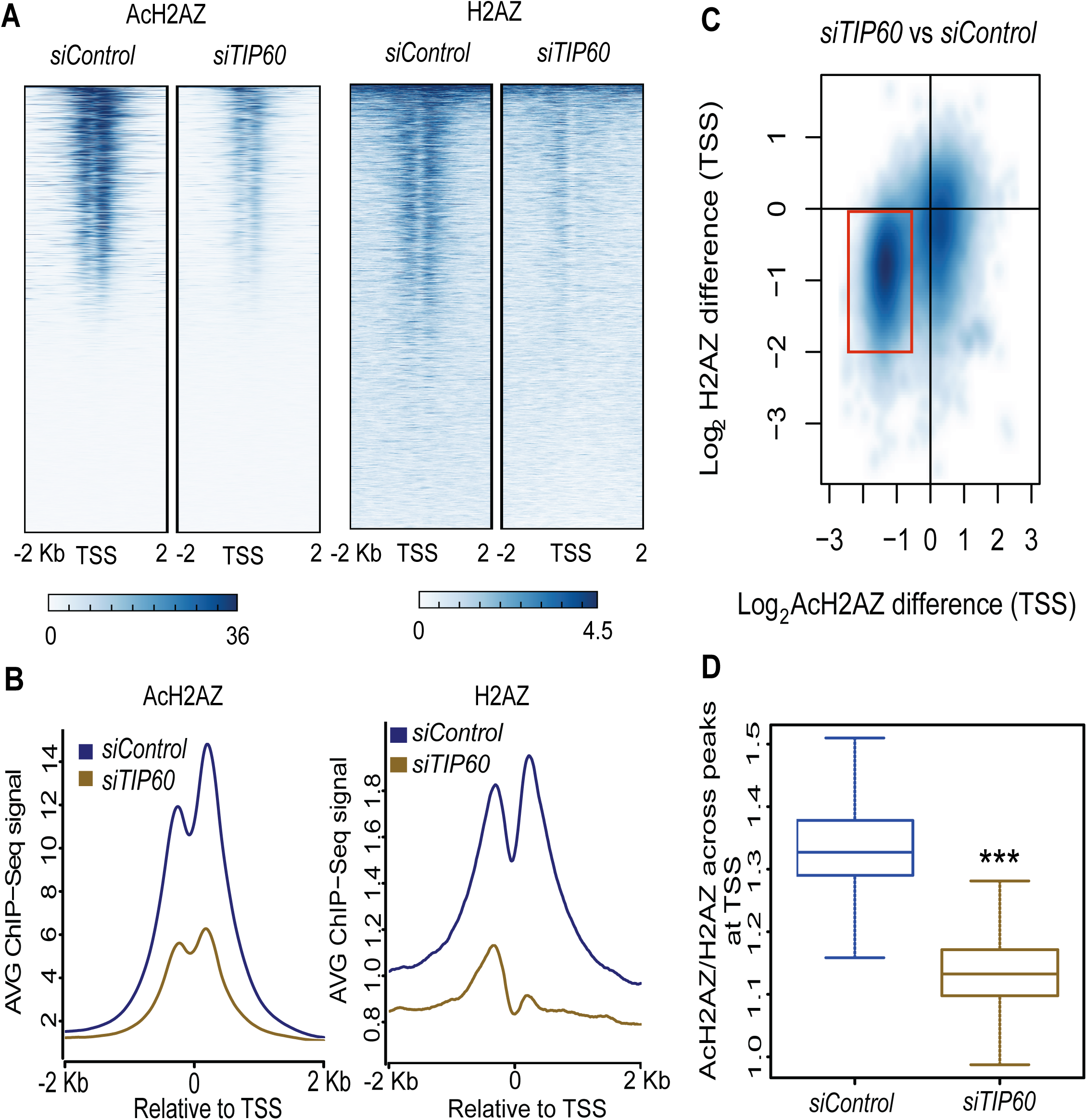
TIP60 regulates AcH2AZ and H2AZ occupancy at the transcription start site (TSS). (**A**) Heatmap displaying the occupancy profiles for AcH2AZ (left) and H2AZ (right) around all the TSS in the human genome (±2 KB) for control and TIP60-depleted MCF10A samples. (N = 1). (**B**) Normalized average signal of the occupancy profiles across all the TSS in the human genome (±2 KB) for AcH2AZ (left) and H2AZ (right) in control (blue) and TIP60-depleted (gold) MCF10A samples. (N = 1). (**C**) Decrease of H2AZ peaks and AcH2AZ peaks on the same locus of the genome. (N = 1). (**D**) Normalization of AcH2AZ occupancy with H2AZ at TSS sites. Boxplot showing the ratio of AcH2AZ/H2AZ at regions surrounding TSS. Differences between the groups were tested using a Wilcoxon signed-rank test. (N = 1) ***, p-value < 0.01.

On comparing the difference of H2AZ and AcH2AZ occupancy across all TSS (+/- 2kb) affected by TIP60 in the genome, we observe a concomitant decrease in occupancy of both H2AZ and AcH2AZ on a large group of TSS’s (Fig. 2C). However, after normalizing AcH2AZ reads to H2AZ, there is still a significant decrease upon depletion of TIP60 (Fig. 2D), suggesting that TIP60 mainly affects H2AZ acetylation at TSS. Taken together, these data suggest that TIP60 mainly regulates the acetylation of H2AZ at TSS, in line with previous findings that TIP60 is involved in the acetylation of H2AZ on chromatin (Altaf et al, 2010; Choi et al, 2009; Dalvai et al, 2013a; Giaimo et al, 2018).

### Transcriptional regulation by TIP60-dependent H2AZ acetylation controls genes involved in DNA repair and cell cycle

Next, we employ RNA-seq to look for the genes with a differential expression upon TIP60 depletion. As shown in Fig. S3A, RNA-seq shows that the depletion of TIP60 results in widespread transcriptional changes. The number of downregulated genes affected by TIP60 depletion is more than that of upregulated genes (Fig. 3A and S3A). Specifically, 523 genes are found significantly upregulated and 850 genes significantly downregulated (|Log_2_ -Fold change| > 1 and FDR < 0.05) (Fig. S3A). To increase the specificity of finding genes whose expression are regulated by TIP60-dependent acetylation of H2AZ, we perform RNA-seq for MCF10A cells transiently depleted of H2AZ. DsiRNA (Kim et al, 2005) which targets both isoforms of H2AZ (siH2AZ) is used, with non-targeting DsiRNA (NC) as control. As shown in Fig. S3B, 525 genes are upregulated and 436 genes are downregulated significantly upon depletion of H2AZ. Pathway analysis from the individual (TIP60 depletion or H2AZ depletion) RNA-seq reveals DNA repair, metabolic processes, and cell cycle-related pathways as common themes (Fig. S4A and S4B).

**Fig. 3.**
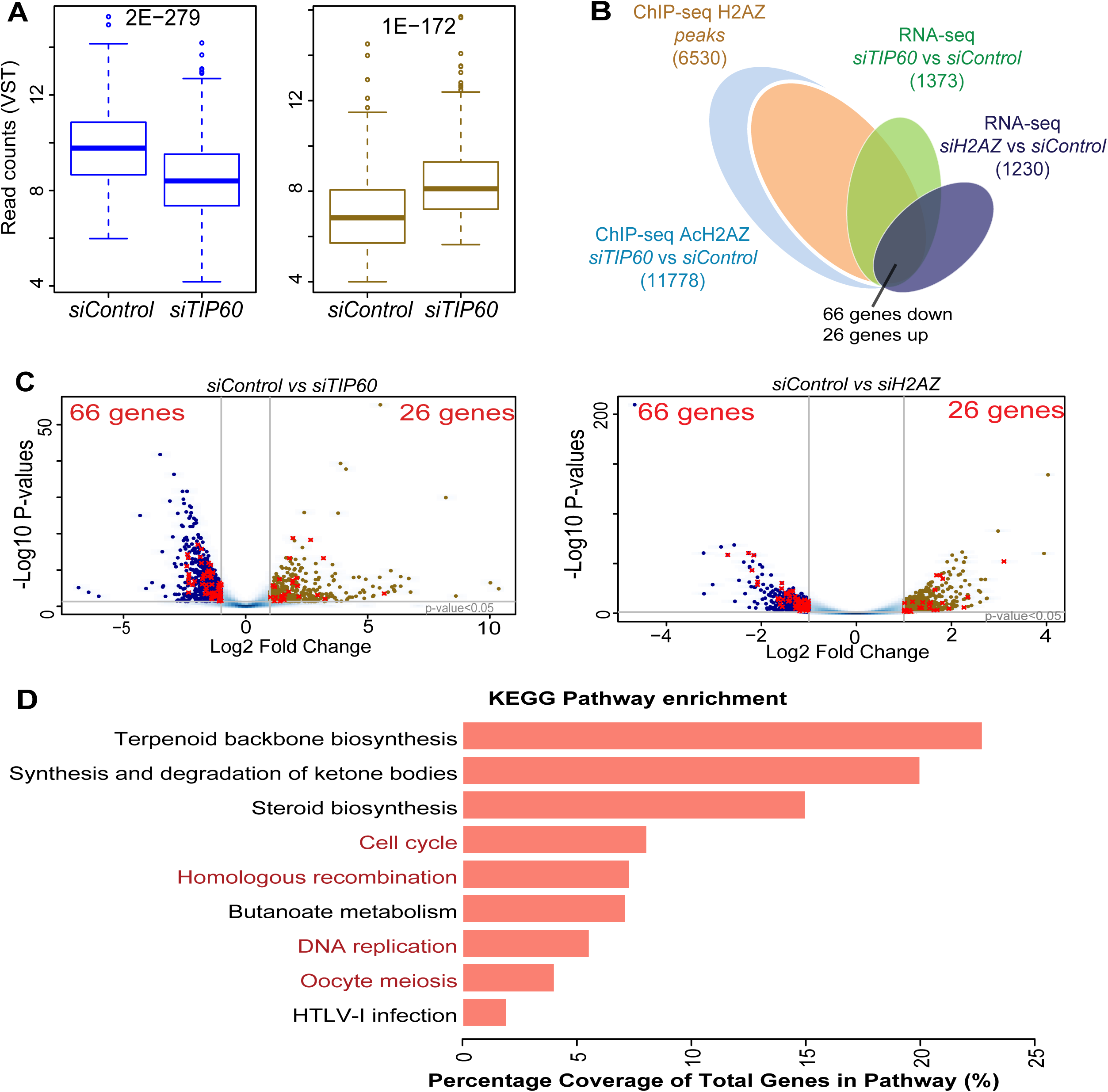
Transcriptional regulation by TIP60-dependent H2AZ acetylation controls genes involved in DNA repair and cell cycle. (**A**) Boxplots represent the expression profile of genes found in downregulated (blue, left) and upregulated (gold, right) groups under control and TIP60-depleted conditions. Values were logarithm transformed using the variance stabilizing transformation (VST). Differences between the groups were tested using a Wilcoxon signed-rank test, the p-value is displayed on top of the boxplots. (N = 2). (**B**) Venn diagram showing the ChIP-seq and RNA-seq overlap strategy. (**C**) Volcano plots showing genes with differential expression after TIP60 (left panel) or H2AZ (right panel) depletion. Dots marked in red represent genes from the overlap shown in B. The overlap selects for genes that have differential occupancy of AcH2AZ and significant occupancy of H2AZ around TSS (±2 KB) between control and TIP60-depleted samples and which show significant expression changes FDR < 0.05 and |log2 (fold change) | > 1 after TIP60 depletion and H2AZ depletion from RNA-seq. Sixty-six downregulated genes and 26 upregulated genes were selected after overlap. (N = 2). (**D**) KEGG pathway enrichment of 66 downregulated genes from the overlap in B. The bars show the percentage of the differentially expressed gene from the total number of genes associated with a given pathway. Pathways related to cell cycle and DNA repair are highlighted in red.

Collectively, to search for differentially expressed genes with TIP60-dependent loss of H2AZ acetylation on their TSS, RNA-seq and ChIP-seq datasets are integrated as shown in the Venn diagram (Fig. 3B). Criteria for selection include a significant decrease in AcH2AZ occupancy on TSS, significant occupancy of H2AZ on TSS, and significant differential expression in TIP60-depleted and H2AZ-depleted RNA-seq datasets. Sixty-six genes from the downregulated group and 26 genes from the upregulated group meet the criteria and are selected for pathway analysis (Fig. 3B and 3C). No significant enriched pathways are found for the upregulated group; however, downregulated genes significantly enriched in biological pathways (Fig. 3D). Notably, out of the multiple pathways regulated by either H2AZ or TIP60 from RNA-seq datasets (Fig. S4A and S4B), cell cycle and homologous recombination (HR) DNA repair pathways are two of the significantly enriched pathways (Fig. 3D). Similar results are also observed in significantly enriched Reactome pathways (Table 1). The 66 downregulated genes used for pathway analysis are shown in Table 2. Since both TIP60 and H2AZ have established roles in DNA repair, we decided to further investigate genes in this pathway to determine if transcriptional regulation of these DNA repair genes could account for their role in DNA repair.

**Table 1.** List of the REACTOME enriched pathways. All pathways are listed with p-value < 0.01.

| **REACTOME Pathway** | **Percent coverage** |
| --- | --- |
| Unwinding of DNA | 41.7 |
| Cholesterol biosynthesis | 33.3 |
| Activation of gene expression by SREBF (SREBP) | 23.1 |
| Regulation of cholesterol biosynthesis by SREBP (SREBF) | 19.4 |
| DNA strand elongation | 15.6 |
| Activation of the pre-replicative complex | 15.6 |
| Assembly of the pre-replicative complex | 15.0 |
| Removal of licensing factors from origins | 14.3 |
| Regulation of DNA replication | 14.3 |
| Activation of ATR in response to replication stress | 13.5 |
| DNA Replication Pre-Initiation | 13.5 |
| M/G1 Transition | 13.5 |
| Polo-like kinase mediated events | 11.8 |
| DNA Replication | 11.7 |
| Orc1 removal from chromatin | 11.5 |
| Switching of origins to a post-replicative state | 11.5 |
| Synthesis of DNA | 10.9 |
| Resolution of D-loop Structures through Synthesis-Dependent Strand Annealing (SDSA) | 10.7 |
| S Phase | 9.8 |
| Inactivation of APC/C via direct inhibition of the APC/C complex | 9.5 |
| Inhibition of the proteolytic activity of APC/C required for the onset of anaphase by mitotic spindle checkpoint components | 9.5 |
| Mitotic Spindle Checkpoint | 9.1 |
| G1/S Transition | 8.6 |
| Resolution of D-loop Structures through Holliday Junction Intermediates | 8.6 |
| Resolution of D-Loop Structures | 8.1 |
| Resolution of Sister Chromatid Cohesion | 7.7 |
| APC-Cdc20 mediated degradation of Nek2A | 7.7 |
| Mitotic G1-G1/S phases | 7.6 |
| Presynaptic phase of homologous DNA pairing and strand exchange | 7.5 |
| Mitotic Anaphase | 7.4 |
| Mitotic Metaphase and Anaphase | 7.4 |
| Separation of Sister Chromatids | 7.3 |
| Cdc20:Phospho-APC/C mediated degradation of Cyclin A | 7.1 |
| Mitotic Prometaphase | 7.1 |
| Homologous DNA Pairing and Strand Exchange | 7.0 |
| APC:Cdc20 mediated degradation of cell cycle proteins prior to satisfaction of the cell cycle checkpoint | 6.9 |
| APC/C:Cdc20 mediated degradation of mitotic proteins | 6.5 |
| Activation of APC/C and APC/C:Cdc20 mediated degradation of mitotic proteins | 6.3 |
| RHO GTPases Activate Formins | 5.9 |
| G2/M Checkpoints | 5.8 |
| Cell Cycle Checkpoints | 5.7 |
| Regulation of APC/C activators between G1/S and early anaphase | 5.6 |
| Cell Cycle, Mitotic | 4.7 |
| HDR through Homologous Recombination (HRR) | 4.3 |
| Cell Cycle | 4.2 |
| M Phase | 3.7 |
| RHO GTPase Effectors | 3.0 |
| Signaling by Rho GTPases | 2.3 |
| Metabolism of lipids and lipoproteins | 1.2 |

**Table 2.**
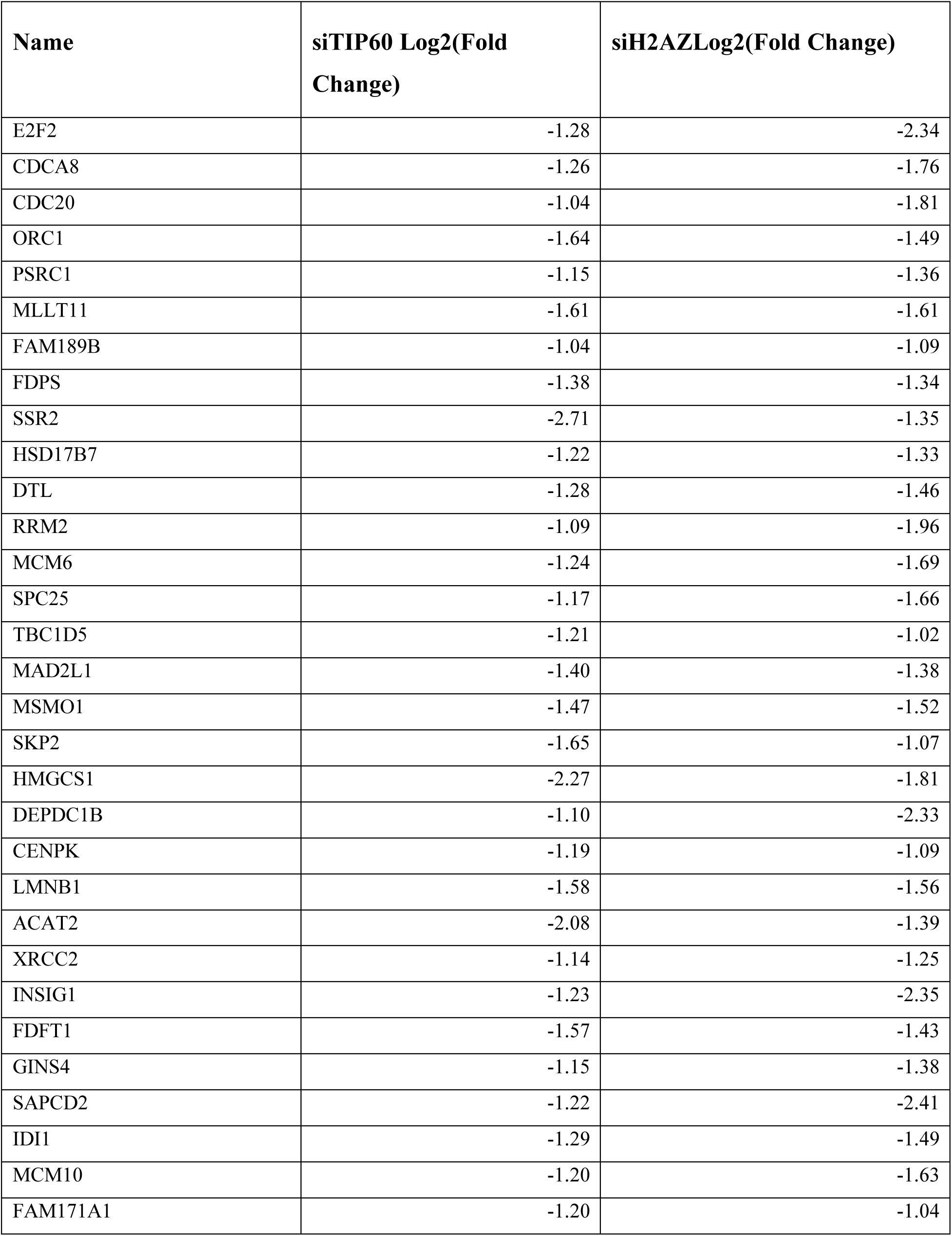
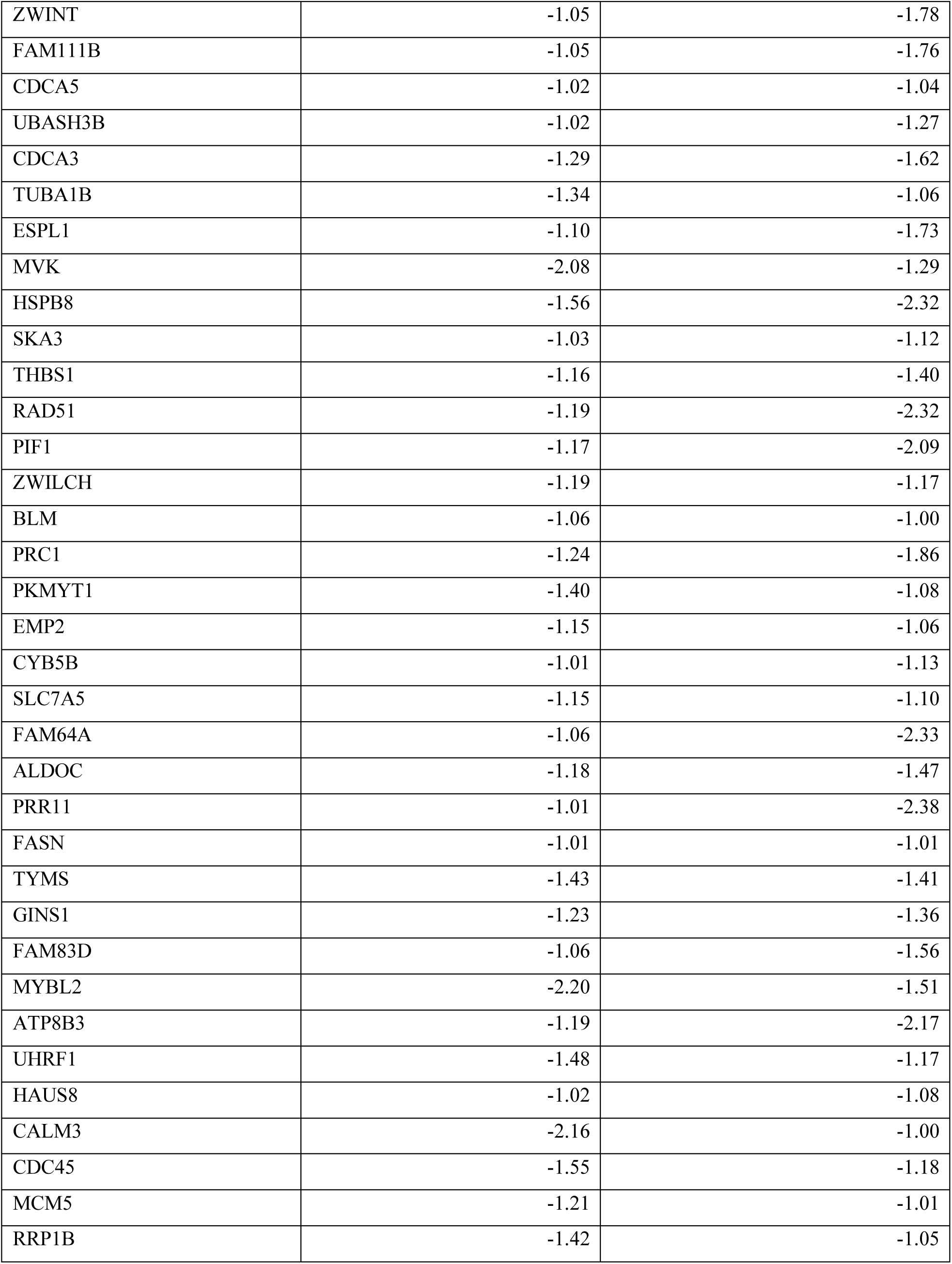
List of genes used in pathway analysis. These genes were downregulated and with a decrease of AcH2AZ occupancy on gene promoter upon TIP60 depletion.

### RAD51 is a DNA repair gene regulated by both TIP60 and H2AZ

Three out of 66 downregulated genes, RAD51 recombinase (*RAD51*), X-ray repair cross-complementing 2 (*XRCC2*) and BLM RecQ like helicase (*BLM*), are enriched in the HR DNA repair pathway. RAD51 is the key protein that replaces Replication protein A1 (RPA1) to bind 3’ single-stranded DNA ends for the homology search and strand invasion steps in HR (Li & Heyer, 2008; Sullivan & Bernstein, 2018). XRCC2 is a paralog of RAD51 which enhances RAD51 foci formation (O’Regan et al, 2001; Sullivan & Bernstein, 2018; Tambini et al, 2010). However, loss of XRCC2 only delays RAD51 foci formation, indicating that XRCC2 is not essential for RAD51 foci formation (Tambini et al, 2010). The role of BLM in HR is unclear (Patel et al, 2017). Studies have suggested that BLM aids in strand resection and in Holliday junction dissolution which promotes HR; however, BLM is also able to inhibit RAD51 filament strand invasion resulting in the creation of “D-loop” structure which inhibits HR (Patel et al, 2017). Given the non-essential function of XRCC2 and the contradictory role of BLM in HR, we choose to focus on RAD51, which has a consistent role in promoting HR DNA damage repair.

Upon depletion of TIP60 or H2AZ in MCF10A, we observe a decrease in both the mRNA and protein levels of RAD51, validating that RAD51 is regulated by TIP60 and H2AZ transcriptionally (Fig. 4A-4D and S5). The change in RAD51 level upon TIP60 depletion is not due to alterations in the cell cycle as we did not observe any cell cycle changes in TIP60-depleted cells (data not shown). *RAD51* mRNA level decrease is also observed in cancer cell lines upon TIP60 depletion (Fig. S6). In addition, we also validate the gene expression of several genes in the cell cycle and DNA repair enriched pathway, which decreases upon TIP60 or H2AZ depletion (Fig. S7A and S7B). This is consistent with a previous study which shows that the mRNA of several HR genes, including *RAD51*, are downregulated upon TIP60 depletion, although the role of H2AZ has not been associated with the process (Su et al, 2017).

**Fig. 4.**
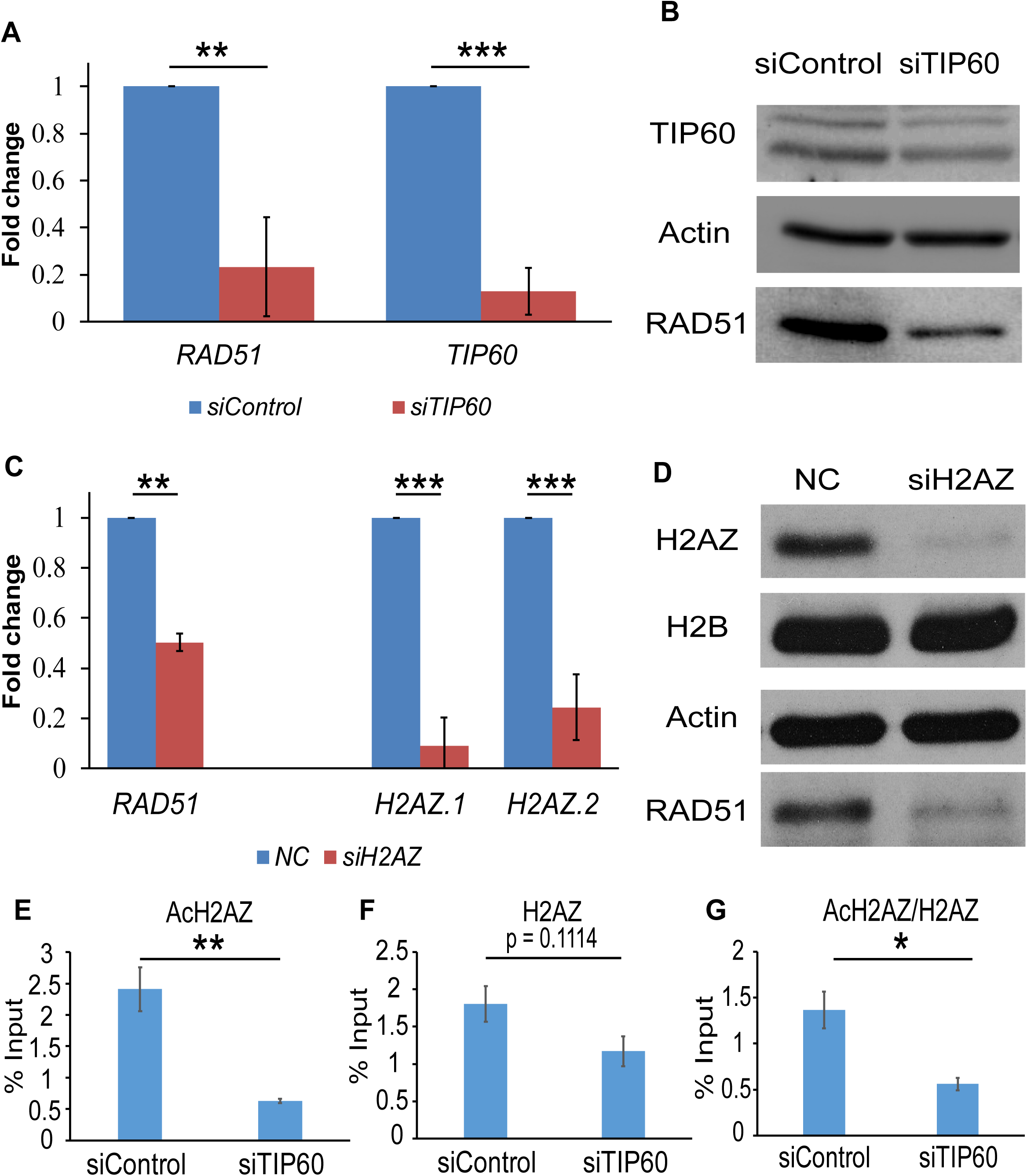
TIP60 and H2AZ regulates RAD51 expression. (**A**) *RAD51* expression upon TIP60 depletion in MCF10A. The expression was validated by qPCR and results were analyzed as fold change against control. All primers used in qPCR experiments are shown in Table 3. (N = 3). All data represent means ± SD. **, p-value < 0.01; ***, p-value < 0.005. (**B**) Western blot showing RAD51 protein level decrease upon TIP60 depletion in MCF10A. Actin serves as a loading control. (N = 2). (**C**) *RAD51* expression upon H2AZ depletion in MCF10A. The expression was validated by qPCR and results were analyzed as fold change against control. All primers used in qPCR experiments are shown in Table 3. (N = 3). All data represent means ± SD. NC, Non-targeting Control. **, p- value < 0.01; ***, p-value < 0.005. (**D**) Western blot showing RAD51 protein level decrease upon H2AZ depletion in MCF10A. Actin and H2B serve as loading controls. (N = 2). (**E-G**) ChIP-qPCR validation showing occupancy of AcH2AZ (E), H2AZ (F), and AcH2AZ/H2AZ ratio (G) on *RAD51* TSS upon TIP60 depletion in MCF10A. (N = 3). All data represent means ± SEM. *, p-value < 0.05; **, p-value < 0.01.

To validate the TIP60-dependent changes in AcH2AZ enrichment on *RAD51* TSS observed in ChIP-seq data, we employ chromatin immunoprecipitation followed by quantitative real-time polymerase chain reaction (ChIP-qPCR). ChIP-qPCR validates the significant decrease in AcH2AZ on the TSS of *RAD51* upon TIP60 depletion (Fig. 4E). A modest but non-significant decrease is also noted for H2AZ occupancy on *RAD51* TSS upon TIP60 depletion (Fig. 4F). Normalizing AcH2AZ to H2AZ signal, TIP60 depletion results in a significant decrease in the AcH2AZ to H2AZ ratio (Fig. 4G), signifying that TIP60 mainly affects acetylation on *RAD51* TSS. In order to exclude the possibility that TIP60 depletion affects nucleosome occupancy on the *RAD51* promoter, we perform H2B ChIP-qPCR. TIP60 depletion does not significantly affect H2B occupancy on *RAD51* TSS (Fig. S8A), indicating that the change in AcH2AZ on *RAD51* TSS is not due to changes in nucleosome occupancy on *RAD51* TSS. A decrease in AcH2AZ to H2B ratio, albeit not significant, is also observed (Fig. S8B).

**Table 3.**
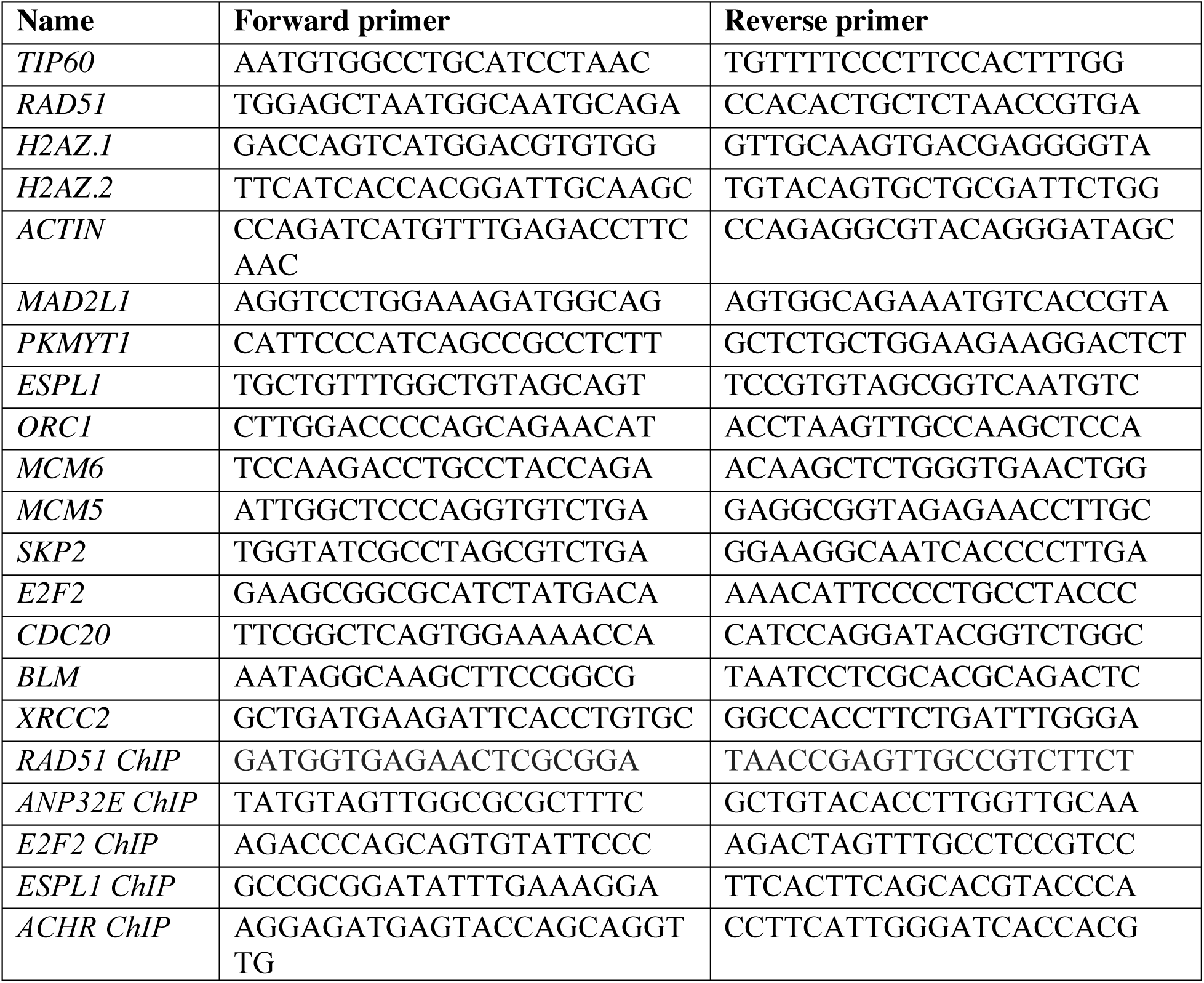
List of qPCR primers used.

AcH2AZ and H2AZ effects on 3 other genes (*ANP32E, ESPL1*, and *E2F2*) and a negative control (*ACHR*) are also validated (Fig. S9). A decrease in AcH2AZ and H2AZ on all 3 genes (*ANP32E, ESPL1*, and *E2F2*), although not significant, is observed while there is no noticeable difference in H2B occupancy (Fig. S9C-9E). Nonetheless, when the AcH2AZ signal is normalized to either H2AZ or H2B, the average % inputs from 3 biological repeats show a significant decrease in the AcH2AZ/H2AZ ratio (for *ANP32E* and *ESPL1*) and AcH2AZ/H2B ratio (for *ANP32E* and *E2F2*) (Fig. S9A and S9B). This validates our ChIP-seq data and indicates that TIP60 regulates mainly the AcH2AZ occupancy and to a lesser extent H2AZ occupancy on the TSS of these genes. In addition, we also performed FLAG-TIP60 ChIP-qPCR on *RAD51* TSS and found that TIP60 enriches on *RAD51* promoter, suggesting a direct regulation (Fig. S10D). This is consistent with other TIP60 ChIP-seq studies in mouse embryonic stem cells (Chen et al, 2015; Ravens et al, 2015) (Fig. S10A and S10B) and FLAG-TIP60 ChIP-seq in human K562 cells (Jacquet et al, 2016) (Fig. S10C), which show TIP60 enrichment on *RAD51* TSS. Taken together, it suggests that TIP60 directly regulates *RAD51* expression mainly through the acetylation of H2AZ on the TSS of *RAD51*.

### TIP60-dependent doxorubicin sensitivity is dependent on the level of RAD51

Multiple reports have shown that the loss of TIP60 increases cellular sensitivity to DNA damage (Hejna et al, 2008; Ikura et al, 2000; Miyamoto et al, 2008; Sun et al, 2005; Tang et al, 2013). We are able to replicate a similar phenotype, where TIP60 depletion sensitizes cells to doxorubicin-induced DNA damage (Fig. 5A).

**Fig. 5.**
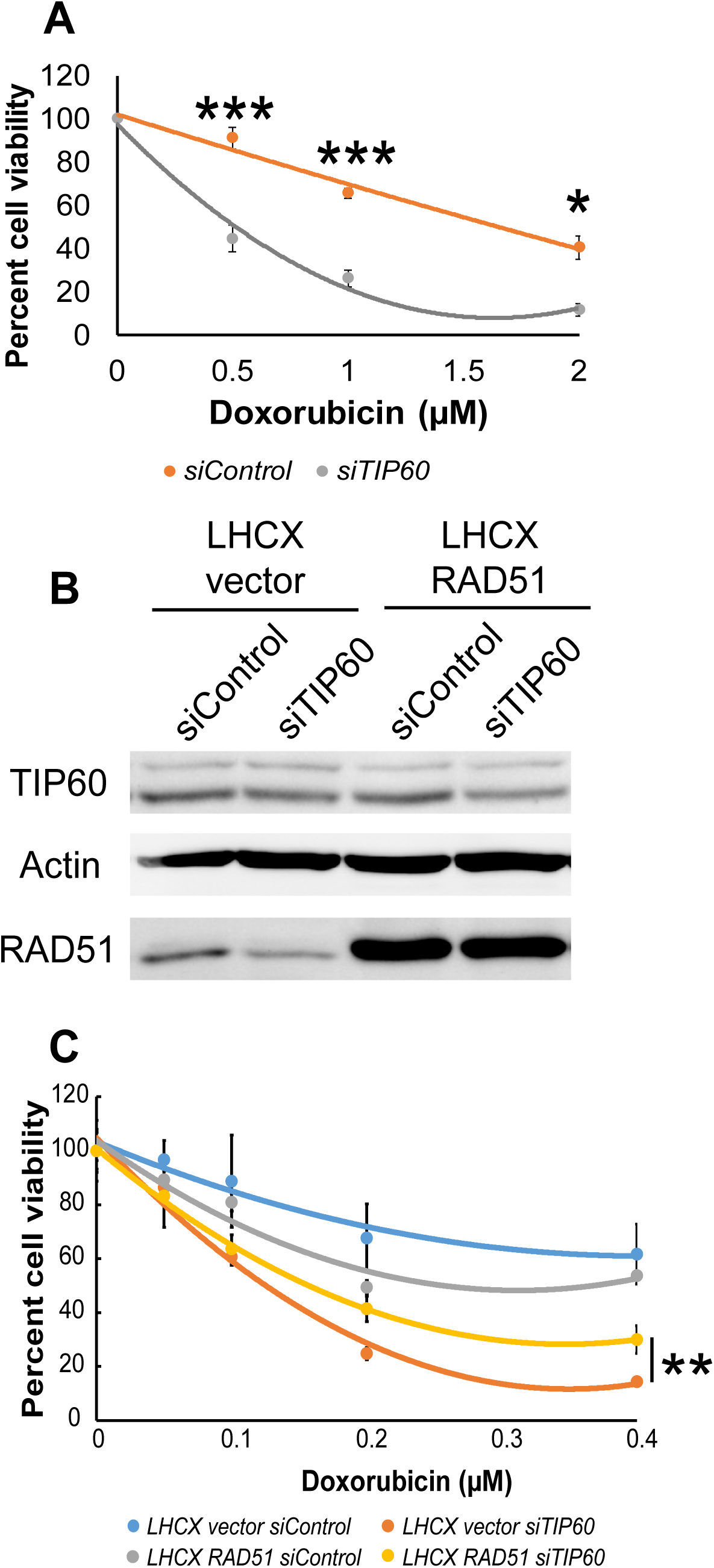
TIP60-dependent doxorubicin sensitivity is partially dependent on the RAD51 level. (**A**) Doxorubicin sensitivity assay showing a decrease in cell viability after depletion of TIP60 and treated with indicated amounts of doxorubicin in MCF10A. Cell viability was measured using an MTS assay. (N = 3). All data represent means ± SEM. *, p-value < 0.05; ***, p-value < 0.005. (**B**) Western blot showing RAD51 protein level changes and expression after TIP60 depletion and RAD51 overexpression in MCF10A. Actin serves as a loading control. (N = 2). (**C**) Representative doxorubicin sensitivity assay showing partial rescue in cell viability in RAD51 expressing MCF10A cells depleted of TIP60 and treated with indicated amounts of doxorubicin from biological repeat 1. Cell viability was measured using an MTS assay. All data represent means ± SD. SD represents the standard deviation from technical replicates. p-value was calculated using technical replicates within the repeat. **, p-value < 0.01.

Studies have shown that TIP60 and H2AZ function mainly through their direct involvement on DNA damage sites (Gursoy-Yuzugullu et al, 2015; Hejna et al, 2008; Ikura et al, 2015; Ikura et al, 2000; Jacquet et al, 2016; Kusch et al, 2004; Squatrito et al, 2006; Sun et al, 2010; Tang et al, 2013; Xu et al, 2012). Two other reports have shown that the expression of several DNA repair genes, such as *ERCC1*, are downregulated upon TIP60 depletion which might contribute to TIP60-dependent DNA damage sensitivity (Miyamoto et al, 2008; Van Den Broeck et al, 2012). Another study found the expression of several DNA repair genes, including *BRCA1, RAD51*, and *FANCD2*, to be regulated by TIP60, and focused on TIP60 regulation on *BRCA1* and *FANCD2* to explain the cisplatin resistance due to TIP60 (Su et al, 2017). In order to determine if regulation of RAD51 level contributes to TIP60-dependent doxorubicin-induced DNA damage sensitivity, we created a stable cell line expressing RAD51 through an exogenous promoter to rescue the phenotype (Fig. 5B). Upon doxorubicin treatment, we observe a consistent trend of partial rescue in the survival of RAD51 expressing cells depleted of TIP60 as compared to vector expressing cells depleted of TIP60, in 3 biological replicates (Fig. 5C and S11). Similar rescue in survival by ectopic RAD51 in doxorubicin-induced DNA damage condition is also observed when cells are depleted of H2AZ, using an siRNA with the same sequence as DsiRNA targeting H2AZ in 2 biological replicates (Fig. S12). Utilizing another DNA damage agent camptothecin, ectopic RAD51 expression is also able to rescue TIP60-dependent camptothecin sensitivity (Fig. S13). However, this is not observed when cells are treated with another DNA damage agent cisplatin (Fig. S14A and S14B), suggesting that the TIP60/H2AZ-regulated *RAD51* expression might be specific to doxorubicin-induced DNA damage. In addition, cisplatin resistance conferred by TIP60 has been previously attributed to TIP60-dependent transcriptional regulation of *ERCC1, BRCA1*, and *FANCD2* (Miyamoto et al, 2008; Su et al, 2017; Van Den Broeck et al, 2012). Overall, these results establish a role for *RAD51* transcriptional regulation in TIP60/H2AZ-dependent doxorubicin-induced DNA damage sensitivity.

## Discussion

In yeast and *Drosophila*, the TIP60 homolog acetylates H2AZ, and aids in the deposition of H2AZ into chromatin (Altaf et al, 2010; Babiarz et al, 2006; Keogh et al, 2006; Krogan et al, 2004; Krogan et al, 2003; Kusch et al, 2004; Mehta et al, 2010; Millar et al, 2006; Mizuguchi et al, 2004). In humans, the dependence of TIP60 for the acetylation of H2AZ on the TSS of some cellular genes has been shown (Dalvai et al, 2013a; Giaimo et al, 2018; Sevilla & Binda, 2014). Additionally, the requirement of TIP60 for the acetylation of H2AZ within octamer or nucleosomes using purified complexes and reconstituted substrates has been established (Choi et al, 2009). TIP60 is also known to exchange H2AZ onto chromatin through the acetylation of H4 and H2A (Altaf et al, 2010; Choi et al, 2009; Kusch et al, 2004) and, possibly through acetylating H2AZ (Giaimo et al, 2018; Millar et al, 2006). There are 2 human isoforms of H2AZ namely, H2AZ.1 and H2AZ.2 (Dryhurst et al, 2009; Horikoshi et al, 2013; Matsuda et al, 2010). Most studies either only focus on H2AZ.1 or do not distinguish between isoforms (Vardabasso et al, 2015). In our study, we show that TIP60 affects H2AZ acetylation in the cellular context (Fig. 1D), in line with previous literature. More importantly, we show that TIP60 does not discriminate between isoforms and can acetylate both isoforms of H2AZ (Fig. 1B and 1C).

ChIP-seq datasets of AcH2AZ and H2AZ upon depletion of TIP60 show loss of both AcH2AZ and H2AZ occupancy on the TSS of cellular genes (Fig. 2), consistent with previous reports that TIP60 is involved in both the exchange and acetylation of H2AZ (Altaf et al, 2010; Choi et al, 2009; Dalvai et al, 2013a; Dalvai et al, 2013b; Giaimo et al, 2018; Keogh et al, 2006; Sevilla & Binda, 2014). Although we achieve low signal intensity from H2AZ ChIP-seq (Fig. 2A and 2B), which could affect the normalization of AcH2AZ to H2AZ signals (Fig. 2D), we have validated *RAD51* and other genes from the ChIP-seq data through ChIP-qPCR (Fig. 4E-4G and S9). Normalizing AcH2AZ to H2AZ in ChIP-qPCR shows a significant decrease upon TIP60 depletion (Fig. 4G, S9A, and S9B), affirming that TIP60 mainly affects H2AZ acetylation on the TSS of *RAD51* and other genes.

Our study is the first to use ChIP-seq to uncover all AcH2AZ and H2AZ occupancy changes genome-wide in a TIP60-depleted condition in human cells (Fig. 2 and S1). Similar genome-wide studies have been done in yeast (Millar et al, 2006), while other reports in human cells are mainly on a single H2AZ regulated gene such as *CCND1* (Dalvai et al, 2013a) or Notch target gene (Giaimo et al, 2018). Several groups also studied global changes in AcH2AZ and H2AZ in human cells but in different contexts such as cancer studies, stem cell differentiation, and in the transcriptional context (Dong et al, 2016; Hardy et al, 2009; Lashgari et al, 2017; Shen et al, 2017; Valdes-Mora et al, 2017; Valdes-Mora et al, 2012; Vardabasso et al, 2015). Nonetheless, when we integrate ChIP-seq data with RNA-seq, the conclusion that AcH2AZ is enriched on actively transcribed genes corroborates with other studies (Dalvai et al, 2013a; Giaimo et al, 2018; Hu et al, 2013; Valdes-Mora et al, 2017; Valdes-Mora et al, 2012). Furthermore, our TIP60-depleted condition strengthens the notion that AcH2AZ and, to a lesser extent, H2AZ occupancy on active gene TSS and gene expression are dependent on TIP60, wherein the loss of TIP60 correlates with loss of AcH2AZ at TSS, modest loss of H2AZ at TSS, and decreased gene expression (Fig. 2-4, S7 and S9).

The TIP60 complex is associated with the acetylation and deposition of H2AZ into chromatin. One possible strategy to study these independent functions of the TIP60 complex and their independent role in transcriptional regulation would be to complement TIP60 depletion with catalytic inactive TIP60 to restore TIP60 complex integrity and H2AZ deposition. However, studies have found that TIP60’s catalytic activity is also involved in H2AZ deposition (Altaf et al, 2010; Choi et al, 2009; Kusch et al, 2004). As such, complementing with a TIP60 catalytic dead mutant will still lead to defects in deposition and total H2AZ occupancy. This highlights the difficulty in separating TIP60’s functions (H2AZ deposition and H2AZ acetylation) in transcriptional regulation.

By combining ChIP-seq and RNA-seq datasets, the expression of 66 genes were identified to be actively regulated through a TIP60/H2AZ-dependent mechanism (Fig. 3B and 3C) (Table 2). The low number of genes (66) from the overlap prompt us to revisit our data. ChIP-seq data shows a large number of peaks (11778) with changes in promoter AcH2AZ occupancy, and 6530 peaks with promoter H2AZ occupancy (Fig. 3B). RNA-seq data upon TIP60 depletion also shows a large number of genes (850 downregulated and 523 upregulated) with changes in gene expression (Fig. S3A). However, RNA-seq with H2AZ depletion shows the least number of gene expression changes (436 downregulated and 525 upregulated) (Fig. S3B), indicating that this dataset is the limiting factor. When the data is re-analyzed without including RNA-seq data for H2AZ depletion, we obtain a resoundingly large number of genes regulated (741 downregulated and 361 upregulated) (Fig. S15), which is almost similar to the number of regulated genes in the RNA-seq data of TIP60 depletion alone (850 downregulated and 523 upregulated) (Fig. 3A), indicating that the majority of genes regulated by TIP60 is through AcH2AZ changes. One possible explanation could be that the H2AZ depletion RNA-seq dataset selects for genes that are most sensitive to changes in H2AZ deposition/level and acetylation, in which both are processes regulated by the TIP60 complex. Therefore, these 66 genes which are the most stringent are subsequently used for pathway analysis.

Pathways enriched from the 66 genes include metabolic pathways (Terpenoid backbone biosynthesis, ketone bodies, steroid biosynthesis, and butanoate metabolism), cell cycle-related pathways (cell cycle, oocyte meiosis, and DNA replication), HR DNA repair pathway, and HTLV-1 infection pathway (Fig. 3D). Interestingly, a recent paper reports that TIP60-dependent lipin-1 acetylation is required for triacylglycerol synthesis in stimulation to fatty acid (Li et al, 2018). Our enrichment of genes involved in metabolic pathways might suggest an additional mechanism in which TIP60 can regulate triacylglycerol synthesis, although a follow-up study needs to be done. Since the role of TIP60 and H2AZ is well-established in DNA damage repair, we decided to explore genes within the HR repair pathway.

Among the genes in the pathway, *RAD51* is validated to be transcriptionally regulated by both TIP60 and H2AZ (Fig. 4). Our finding that H2AZ regulates *RAD51* is supported by another study where microarray analysis is performed for cells depleted specifically of either H2AZ.1 or H2AZ.2 to look for global gene changes, and integrated with H2AZ ChIP-seq (Vardabasso et al, 2015). While genes involved in immunological pathways are enriched for H2AZ.1 depletion, H2AZ.2 depletion enriched for genes involved in the cell cycle (Vardabasso et al, 2015), similar to what we observe in our datasets (Fig. 3D). Furthermore, *RAD51* and several other genes including, *ORC1, MCM5, MCM6, ESPL1, XRCC2*, and *BLM* are found to be among the genes regulated by H2AZ.2 but not H2AZ.1 (shown in the supplementary data of their study) (Vardabasso et al, 2015) and are consistent with our data that H2AZ regulates these genes (Fig. 4C, 4D, and S7B). In addition, this implies that the TIP60-dependent transcriptional regulation of *RAD51* might be dependent on H2AZ.2. Indeed, when we use siRNA specific for individual H2AZ isoforms, H2AZ.2 depletion seems to correlate more with the decrease in the regulated genes (Fig. S16). Taken together with our data that TIP60 acetylates H2AZ.2 (Fig. 1C) and that AcH2AZ occupancy is dependent on TIP60 (Fig. 4E-4G), it suggests that TIP60 might regulate *RAD51* transcriptionally through H2AZ.2 acetylation on TSS.

Apart from validating AcH2AZ and H2AZ changes on *RAD51*, 3 other genes (*ANP32E, ESPL1*, and *E2F2*) are also validated from the ChIP-seq data (Fig. S9). Interestingly, ANP32E the histone chaperone which removes H2AZ from chromatin, is regulated positively by TIP60 which is involved in H2AZ deposition. This might suggest a potential feedback mechanism in cells to try to maintain the equilibrium of chromatin H2AZ.

In this study, we found that TIP60 localizes to *RAD51* TSS (Fig. S10D). A key question that remains to be addressed is how TIP60 is recruited to the TSS of *RAD51*. To address this, we performed a preliminary bioinformatics search for transcription factors that can regulate the 66 down-regulated genes identified using the Transcriptional Regulatory Relationships Unraveled by Sentence-based Text mining (TRRUST) tool (Han et al, 2018). Among the transcription factors predicted to regulate these genes, only E2F1 and E2F4 are predicted to regulate *RAD51* (Fig. S17). E2F1 interacts with and recruits TIP60 to promoters of cell cycle genes to regulate their expression (Taubert et al, 2004) as such, E2F1 could be the transcription factor that recruits TIP60 to H2AZ regulated sites. However, it has also been shown that E2F1 recruitment is dependent on H2AZ.2 (Vardabasso et al, 2015), indicating that E2F1 would probably be downstream of TIP60 since TIP60 regulates the acetylation and deposition of H2AZ (Fig. 2). In addition, TIP60 is known to acetylate E2F1 to promote its loading on chromatin, supporting that E2F1 is downstream of TIP60’s activity (Van Den Broeck et al, 2012). There are currently no reports on E2F4 and TIP60, and E2F4 is known to repress *RAD51* expression during hypoxia and is dependent on its complex formation with p130 (Bindra & Glazer, 2006; Hegan et al, 2010); therefore, it will be interesting if the repressor E2F4 can recruit TIP60 to regulate acetylation of H2AZ for transcription activation in non-hypoxic conditions.

H2AZ containing nucleosomes are less stable than H2A containing nucleosomes (Jin & Felsenfeld, 2007; Jin et al, 2009; Zhang et al, 2005). As such, the deposition of H2AZ into *RAD51* TSS by TIP60 could lead to an open conformation, permitting the recruitment of transcription factors and transcription activation. With regard to acetylation, H2AZ acetylation with core histone acetylation synergizes and destabilizes nucleosomes (Ishibashi et al, 2009). Since TIP60 can acetylate several core histones and H2AZ (Kimura & Horikoshi, 1998; Yamamoto & Horikoshi, 1997) (Fig. 1), it is likely that the mechanism of TIP60-mediated *RAD51* transcriptional activation could be through nucleosome destabilization, and requires TIP60’s acetylation on both core histones and H2AZ. Another possible mechanism is through the recruitment of transcriptional activator BRD2. Attempts to discover downstream effector proteins that are recruited by H2AZ reveal that BRD2 is recruited by a combination of H2AZ and acetylated H4 on androgen receptor-regulated genes (Draker et al, 2012). Moreover, BRD2 recruitment on cell cycle genes and subsequent transcriptional activation is dependent on H2AZ.2 (Vardabasso et al, 2015).

Interestingly, we observe that *H2AZ* expression decreases upon TIP60 depletion albeit with variable fold changes (Fig. S18A and S18B), suggesting that TIP60 might regulate H2AZ transcriptionally, which in turn regulates *RAD51* expression. This could be through a plausible self-regulatory mechanism of H2AZ as observed from a decrease in AcH2AZ and H2AZ promoter occupancy on *H2AFZ* and *H2AFV* upon TIP60 depletion from our ChIP-seq data (Fig. S18C and S18D). Taken together, it posits an additional mechanism in which TIP60 could be acting to regulate *RAD51* through regulating *H2AZ* transcriptionally, although a follow-up study will be required.

Both TIP60 and H2AZ play crucial roles in DNA damage repair through their recruitment to DNA damage sites (Gursoy-Yuzugullu et al, 2015; Hejna et al, 2008; Ikura et al, 2015; Ikura et al, 2000; Jacquet et al, 2016; Kusch et al, 2004; Squatrito et al, 2006; Sun et al, 2010; Tang et al, 2013; Xu et al, 2012). To our knowledge, three reports have shown that the expression of several DNA repair genes are dependent on TIP60, and could partially explain TIP60-dependent cellular sensitivity to cisplatin-induced DNA damage (Miyamoto et al, 2008; Su et al, 2017; Van Den Broeck et al, 2012). However, the authors focused on *BRCA1, FANCD2*, and *ERCC2* to explain the chemoresistance of cisplatin due to TIP60 (Su et al, 2017; Van Den Broeck et al, 2012). In addition, the role of H2AZ in the regulation of these DNA repair genes, and the contribution of *RAD51* transcription regulation to TIP60-dependent DNA damage sensitivity have not been established. Our data where TIP60/H2AZ-dependent doxorubicin, but not cisplatin, sensitivity is rescued by restoring the level of RAD51 (Fig. 5C, S11 and S12) delineates a role for *RAD51* transcription in TIP60/H2AZ-dependent doxorubicin-induced DNA damage sensitivity. In addition, ectopic expression of H2AZ to rescue the H2AZ-dependent sensitivity could be done to show the specificity of H2AZ in this phenomenon. Even though we show that *RAD51* is regulated by TIP60 in several cell lines (Fig. S6), doxorubicin DNA damage sensitivity assays could be done in these cell lines to show generality between cell lines.

The decrease in *RAD51* upon TIP60 depletion forms at least partly the basis of the increased cellular sensitivity observed in TIP60-depleted cells to doxorubicin Anthracyclines form the cornerstone of chemotherapy in breast cancer. In this context, the identification of pathways that confer anthracycline sensitivity is vital. Our study in MCF10A suggests that modulation of TIP60 and H2AZ potentiates anthracycline sensitivity in a manner dependent on RAD51. In line with these findings, an inhibitor of RAD51 has been shown to sensitize multiple myeloma cells to doxorubicin, indicating the importance of RAD51 level to cellular survival in response to doxorubicin treatment (Alagpulinsa et al, 2014). In conclusion, our study shows that TIP60 and H2AZ regulate *RAD51* expression and specifies a TIP60/H2AZ-dependent transcriptional role in doxorubicin-induced DNA damage repair (Fig. 6). These findings could be expanded upon in the future to study the correlation of TIP60 dysregulation in breast cancer with chemotherapy outcomes, and also evaluate small molecules targeting this pathway as potential chemosensitizers.

**Fig. 6.**
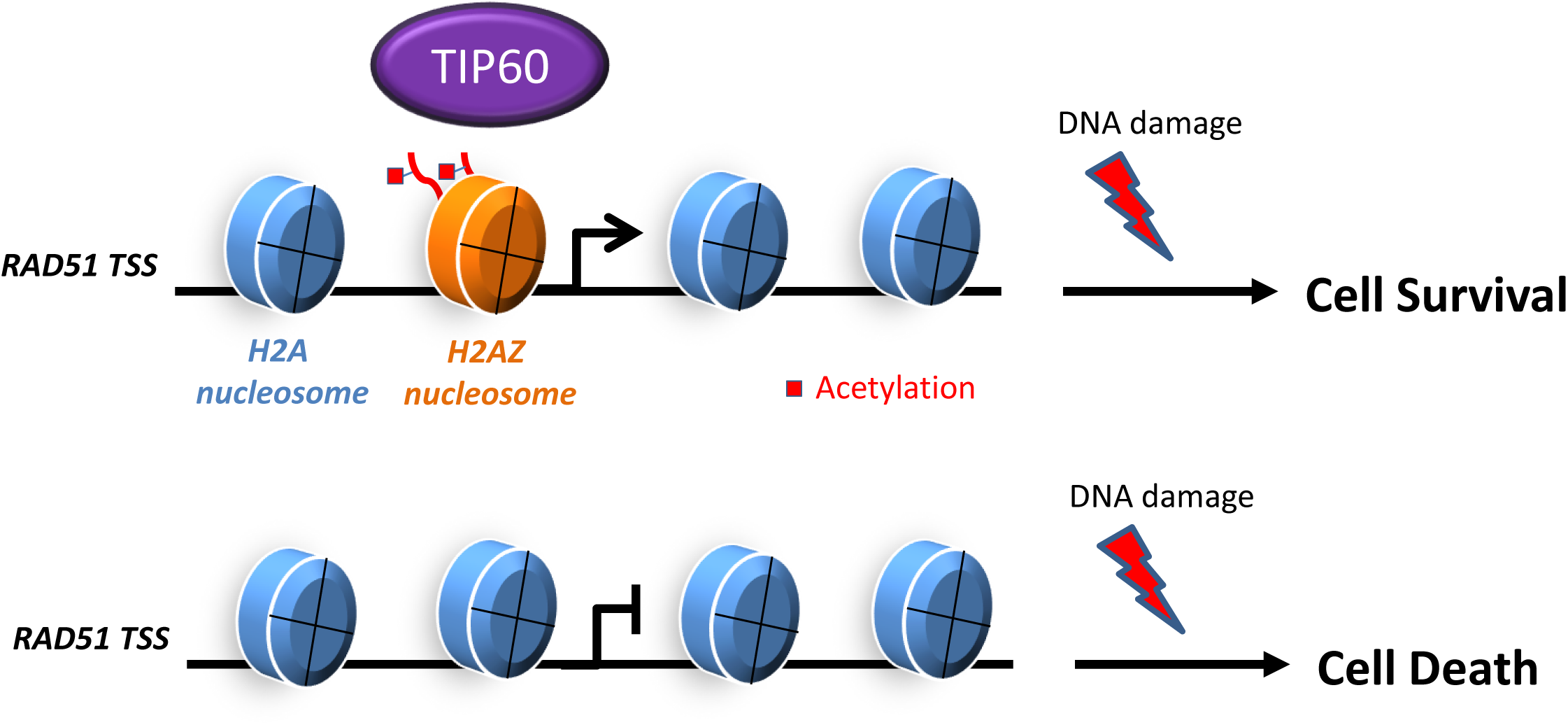
Schematic model of TIP60 regulation on *RAD51* TSS and cellular outcome upon DNA damage. TIP60 mediates the acetylation of H2AZ on *RAD51* TSS, resulting in *RAD51* gene expression and cell survival after DNA damage. In the absence of TIP60, *RAD51* expression decreases which leads to cell death upon DNA damage.

## Materials and Methods

### Cloning

H2AZ.1 and H2AZ.2 were cloned into pET28A(+) vector using *Bam*HI and *Not*I restriction sites. *RAD51* was cloned into pLHCX-hygro vector using *Not*I site, introduced using *Hind*III site. All plasmids were confirmed through sequencing.

### Bacterial recombinant protein purification

pET28a-His-TIP60, pET28a-His-H2AZ.1, and pET28a-His-H2AZ.2 constructs were transformed into BL21 (DE3) competent cells through pre-cooling on ice for 30 min and subsequent heat shock at 42°C for 30 sec. The cells were cooled on ice for 5 min before plated on kanamycin agar plates and incubated overnight at 37°C. A bacterial colony was picked and inoculated into LB for growth at 37°C until 0.5 optical density (OD) of the culture was reached. The expression of recombinant protein (rTIP60, rH2AZ.1, and rH2AZ.2) was induced by treating the culture with isopropyl β-D-1-thiogalactopyranoside (IPTG) (0.5 mM final concentration) for 16 h at 18°C (rTIP60) or 3 h at 37°C (rH2AZ.1 and rH2AZ.2). Bacterial cells were harvested and suspended in bacterial lysis buffer (50 mM Tris-HCl, pH 8.0, 150 mM NaCl, 10 mM Imidazole, 1 mM PMSF, Protease inhibitor cocktail, and 10% Glycerol). Lysozyme (10 mg/ml) was added and incubated at 37°C for 30 min, followed by a 6-cycle sonication at 25% amplitude with 10 sec on and 30 sec off. Triton-X-100 (final conc. 0.1%) (Sigma-Aldrich, Cat. No. T8787) was added and rotated at 4°C for 1 h. The soluble fraction and the pellet were separated by spinning at 13,000 rpm for 15 min at 4°C.

For recombinant TIP60 (rTIP60) protein, the supernatant was added to 2 ml of pre-equilibrated Nickel-nitrilotriacetic acid (Ni-NTA) agarose (QIAGEN, Cat. No. 30210) 50% slurry and rotated at 4°C for 2 h. For recombinant H2AZ (rH2AZ.1 and rH2AZ.2) protein, the pellet was dissolved in 2 ml Dimethyl sulfoxide (DMSO) (MP BIOMEDICALS, Cat. No. 02191418) by stirring using a magnetic stirrer for 30 min at room temperature. Ten milliliters of unfolding buffer [bacterial lysis buffer with 6 M Guanidinium-HCl (Amresco, Cat. No. 50-01-1)] was added to the mixture and rotated at 37°C for 1 h, followed by centrifugation at room temperature for 30 min at 12,000 rpm. The supernatant was collected and incubated with 2 ml Ni-NTA agarose 50% slurry at 4°C for 2 h.

The Ni-NTA agarose and bounded proteins were washed three times with wash buffer B (50 mM Tris-HCl pH 8.0, 150 mM NaCl, 40 mM Imidazole, 10% Glycerol, and 6 M Guanidinium-HCl was added for H2AZ.1 and H2AZ.2 purification) by rotating at 4°C for 5 min each. Proteins were eluted in elution buffer (50 mM Tris-HCl pH 8.0, 150 mM NaCl, 250 mM Imidazole, 10% Glycerol, and 6M Guanidinium-HCl was added for H2AZ.1 and H2AZ.2 purification) by rotating at 4°C for 30 min. The eluted proteins were dialyzed overnight at 4°C against HAT assay buffer (50 mM Tris, pH 8.0, 0.1 mM EDTA, 1 mM DTT, and 10% Glycerol).

### In vitro HAT assay

Purified recombinant TIP60 (0.5 - 2 µg) was incubated in 40 µl reaction mixture containing 2 µg of purified recombinant H2AZ.1 or H2AZ.2, 100 µM acetyl-coenzyme A (AcCo-A) and HAT assay buffer at 30°C for 10 min. To stop the reaction, 40 µl of SDS loading dye (2×) was added. Samples were boiled in 95°C for 5 min before loading into 15% SDS-PAGE gel.

### Cell culture

MCF10A cells (ATCC® CRL-10317™) were cultured in Dulbecco’s modified Eagle’s media (DMEM)/F12 (1:1) media (Gibco, Cat. No. 11330-032) with 5% horse serum (Gibco, Cat.No. 16050-122), 1% penicillin-streptomycin (Gibco, Cat.No. 15140-122), 20 ng/ml epithelial growth factor (Peprotech, Cat.No.AF-100-15), 0.5 mg/ml hydrocortisone (Sigma-Aldrich, Cat. No. H-0888), 100 ng/ml cholera toxin (Sigma-Aldrich, Cat. No. C-8052) and 10 µg/ml insulin (Sigma-Aldrich, Cat. No. I-1882). 293T (ATCC® CRL-3216™), HEPG2 (ATCC® HB-8065™) and HCT116 (ATCC® CCL-247™) cells were cultured in Dulbecco’s modified Eagle’s media (DMEM) high glucose (Sigma, Cat. No. D-5796) with 10% fetal bovine serum (Sigma, Cat. No. F-7524), 1% penicillin-streptomycin (Gibco, Cat.No. 15140-122). DLD1 (ATCC® CCL-221™) and SNU398 (ATCC® CRL-2233™) cells were grown in RPMI 1640 Media (HyClone Cat. No. SH30027.01) with 10% fetal bovine serum (Sigma, Cat. No. F-7524), 1% penicillin-streptomycin (Gibco, Cat.No. 15140-122). Cells were incubated at 37°C with 5% CO_2_.

### Stable cell lines

Virus was generated by transfecting 5×10^6^ 293T cells with the plasmids MSCV construct: i.e., MSCV vector control, TIP60 wild-type (TIP60WT), siRNA-resistant TIP60 wild-type (TIP60*WT), siRNA-resistant catalytically inactive TIP60 (TIP60*KD) using Lipofectamine 2000 (Invitrogen/Life Technologies, Cat. No. 52887), as per manufacturer’s protocol. Viruses were harvested after 72 h of transfection and were used to infect 1×10^6^ MCF10A together with polybrene (Sigma-Aldrich, Cat. No. S2667) reagent (4 µg/ml). After 6 h, media containing the virus was replaced by growth media. After 24 h, puromycin (1 µg/ml) was added into the growth media for selection. Media with antibiotics was changed every 48 h until the mock-transfected cells died. The cells were continuously selected for 2 weeks and for the creation of stable cell lines. The same protocol was used for the generation of MCF10A LHCX vector and LHCX-RAD51 stable cell lines using hygromycin (50 µg/ml) for selection.

### siRNA sequences

siControl, siTIP60, siH2AZ.1, and siH2AZ.2, siH2AZ (siRNA) were purchased from Sigma Aldrich. siH2AZ and negative control (NC) were DsiRNA from Integrated DNA Technologies. siTIP60-2 is a commercially available siRNA purchased from Sigma Aldrich (Cat. No. SASI_Hs01_00073301).

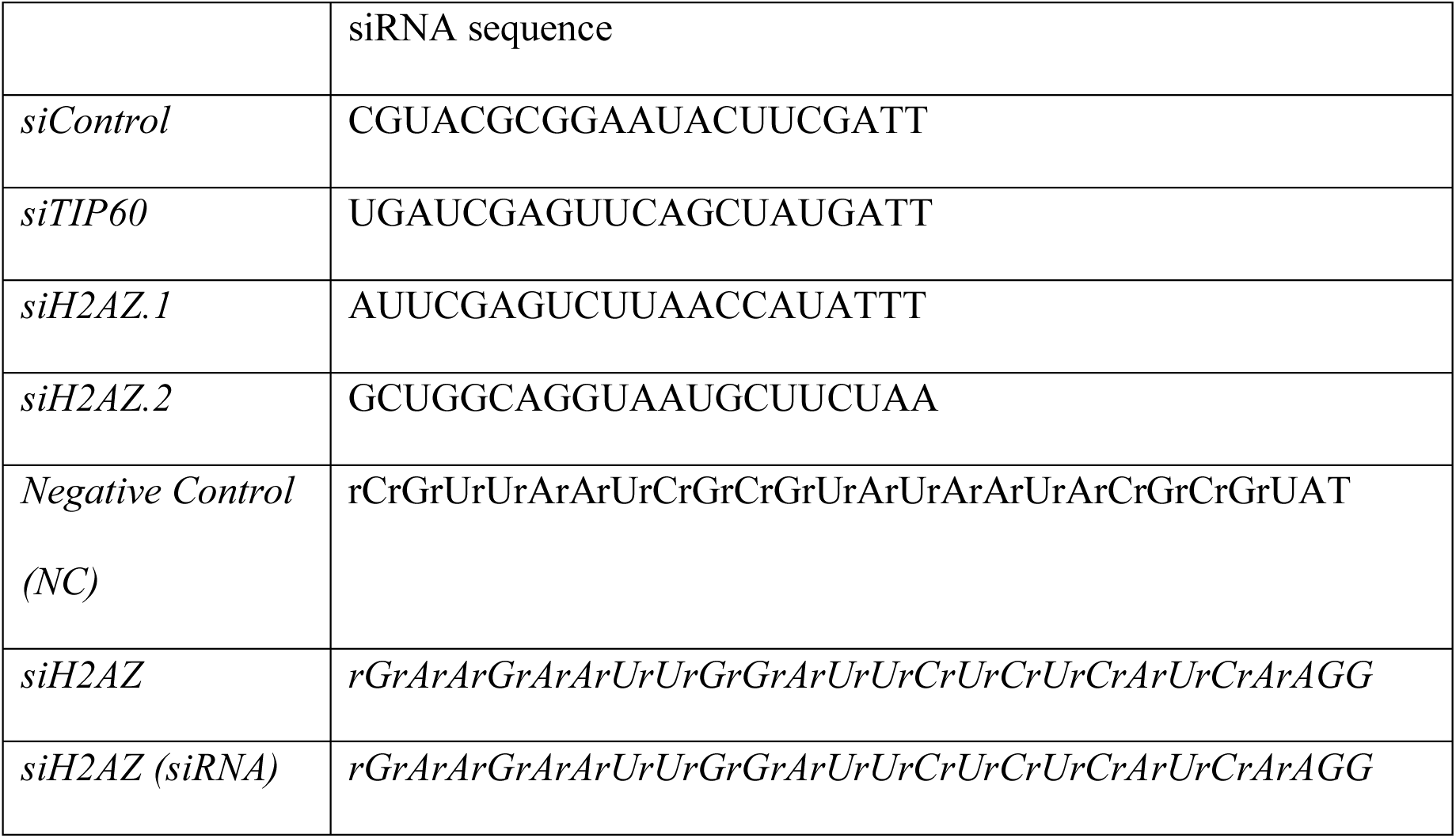

### siRNA transfection

siRNAs targeting different genes were transfected using Lipofectamine RNAiMax reagent from Invitrogen/Life Technologies (Cat. No. 56532). The siRNA mixture used contained 15 μl Lipofectamine RNAiMax, 2 ml Opti-MEM media (Gibco, Cat. No. 31985-070) and 10 nM siRNA for MCF10A transfection and 20 nM for transfecting other cell lines. The transfection mixture was incubated for 20 min at room temperature and then 1×10^6^ cells were seeded in a 10-cm plate together with the siRNA mixture in 5 ml. For MCF10A, the transfection mixture was replaced by growth media after 6 h. Twenty-four hours later from the start of transfection, the second round of knockdown using the same protocol was performed. Cells were harvested 72 h from the first round of siRNA transfection. For other cell lines, transfection mixture was replaced by growth media after 24 h, and cells harvested after 72 h.

### Purification of total RNA

Total RNA was isolated using TRIZOL reagent (Invitrogen/Life Technologies, Cat No. 15596-026) following the manufacturer’s protocol. RNA was dissolved in nuclease-free water for reverse transcription-PCR.

### Reverse transcription and quantitative PCR

Complementary DNA (cDNA) was synthesized using the iScript cDNA Synthesis Kit (Bio-Rad, Cat. No. 170-8891) according to the manufacturer’s protocols. Quantitative PCR (qPCR) was performed with primer sets corresponding to primer list (Table 3) and using iTaq Universal SYBR Green Supermix (Bio-Rad, Cat. No. 172-5125) on an Applied Biosystems 7500 Fast Real-Time PCR system or ThermoFisher Scientific QuantStudio 3 or 5 Real-Time PCR system. Results were analyzed by calculating ΔΔCT and are represented as fold change. Actin was used as the control gene.

### Western blotting

Cells were lysed in lysis buffer (10 mM Tris-HCl pH 7.5, 150 mM NaCl, 0.5% NP-40, and 0.5 mM EDTA) with protease inhibitors (Nacalai Tesque, Cat. No. 04080-11). Proteins were separated on SDS-PAGE gel, transferred onto a nitrocellulose membrane (BioRad, Cat. No. 9004-70-0) and detected using indicated antibodies.

### Antibodies

Antibodies used were the following. β-actin (SantaCruz Biotechnology, Cat.No. sc-81178, 1:1000); H2AZ (Abcam, Cat. No. ab4174, 1:2000); AcH2AZ (Abcam, Cat.No. ab18262, 1:2000); H2B (Abcam, Cat. No. ab1790, 1:2000); and RAD51 [14B4] (GeneTex, Cat. No. GTX70230, 1:250) or RAD51 (Abcam, Cat No. ab133534, 1:10,000). TIP60 rabbit polyclonal antibody was generated in the lab and reported previously (Frank et al, 2003; Jha et al, 2010; Rajagopalan et al, 2017; Rajagopalan et al, 2018; Subbaiah et al, 2016; Zhang et al, 2016).

### Chromatin immunoprecipitation (ChIP)

ChIP was performed as described earlier (Jha et al, 2010; Karnani et al, 2007). Briefly, 1.5×10^6^ cells were transfected with siControl or siTIP60 in a 10-cm plate. After 48 h of siRNA treatment, cells were cross-linked with 1% formaldehyde (SantaCruz Biotechnology, Cat. No. sc-203049A) for 10 min at room temperature, and then washed twice with ice-cold phosphate-buffered saline. Cells were harvested by scraping and centrifuged at 1750 g for 15 min to collect the cell pellet. Cells were then resuspended in SDS lysis buffer (1% SDS, 0.01 M EDTA, and 0.05 M Tris-HCl, pH 8.0) and sonicated (ON: 15 sec and OFF: 45 sec at 30% amplitude for 40 cycles) to obtain DNA fragments ranging from 100 to 500 bps. Chromatin was isolated by centrifuging at 15300 g for 15 min at 4°C and the supernatant was collected for immunoprecipitation.

Immunoprecipitation was performed by incubating anti-H2AZ and anti-AcH2AZ antibody with protein A/G PLUS-agarose beads (SantaCruz Biotechnology, Cat. No. sc-2003) overnight at 4°C. Beads were then washed with (i) low-salt immune complex buffer (0.1% SDS, 1% TritonX-100, 0.002 M EDTA, 0.02 M Tris-HCl, pH 8.0, and 0.15 M NaCl), (ii) high-salt immune complex buffer (0.1% SDS, 1% Triton-X-100, 0.002 M EDTA, 0.02 M Tris-HCl, pH 8.0, and 0.5 M NaCl), (iii) LiCl buffer (0.25 M lithium chloride, 1% NP40, 0.005 M EDTA, 0.01 M Tris-HCl, pH 8.0, and 1% deoxycholate) and (iv) TE buffer (0.005 M EDTA and 0.01 M Tris-HCl, pH 8.0). Beads were eluted in 100 µl elution buffer (1% SDS and 0.0084% NaHCO_3_) three times with agitation for 15 min each. Chromatin was reversed cross-linked by adding 0.2 M NaCl and was heated at 65°C for overnight. The proteins bound to DNA were digested by adding 20 µg proteinase K (AppliChem, Cat. No. 39450-01-6) and incubated at 45°C for 1 h. DNA was purified using a PCR purification kit (QIAGEN, Cat. No. 28106).

### DNA damage sensitivity assay

To evaluate the effects of doxorubicin/cisplatin/camptothecin on cell viability of MCF10A upon depletion of TIP60, 3-(4,5-dimethylthiazol-2-yl)-5-(3-carboxymethoxyphenyl)-2-(4-sulfophenyl)-2H-tetrazolium (MTS) (Promega, Cat. No. G109C) cell viability assay was performed. Cells were seeded in triplicates at 1×10^4^ cells/well in a 96-well plate containing a final volume of 100 µl/well. Each sample was exposed to vehicle control or DNA damage agent at different concentrations. Vehicle control for doxorubicin and campothecin was DMSO and vehicle control for cisplatin was DMF. Cells were treated with doxorubicin or camptothecin for 24 h, while cells were treated with cisplatin for 96 h. After the indicated period of exposure, 20 µl of MTS solution was added to each well to achieve a final concentration of 0.33 mg/ml. The plate was incubated at 37°C and the absorbance of each well was taken at 490 nm at each hour for a total of 4 h. The background absorbance in each well was normalized by subtracting the absorbance value of a well without cells to increase the signal-to-noise ratio. The absorbance obtained from doxorubicin-exposed wells is presented as a percentage of the normalized absorbance of the vehicle control (DMSO/DMF).

### ChIP-seq sequence alignment

ChIP-seq reads were aligned to the human reference genome GRCh37/hg19 using ‘STAR Aligner’ (v2.5.1b) (Dobin et al, 2013) with EndToEnd as the alignEndsType option and 1 as the alignIntronMax threshold. All other parameters were set to the default. Read duplicates were removed from the bam files using the ‘MarkDuplicates’ module from ‘Picard’ (v1.140). BigWig files were generated from the resulting bam files using the ‘bamCoverage’ from ‘Deeptools’ (v2.2.2-1) (Ramirez et al, 2016), reads were extended 250 base-pairs (bp) and normalized per genomic content as follows: number of reads / ((total number of mapped reads * fragment length) / effective genome size), where the effective genome size is 2,451,960,000 base-pairs. Bigwig files were loaded into ‘IGV’ (v) for visualization.

### ChIP-seq post-alignment processing

Differential binding analysis and the annotation of the differentially bound regions was performed with ‘diffReps’ (v1.55.6) (Shen et al, 2013) using the G-test method with 300bp of shift and all other parameters were set to the default. Bam files without duplicated reads were converted to bed format using the ‘bamtobed’ script from ‘bedtools’ (v2.25.0) (Quinlan & Hall, 2010). Heatmaps and average plots were generated by ‘computeMatrix’ (reference-point mode) with default parameters and ‘plotHeatmap’ from the ‘deeptools’ (v2.2.2) suite (Ramirez et al, 2016). All scatterplots describing the occupancy differences were generated as following: ‘ComputeMatrix’ was used to count the normalized occupancy signal from the bigwig files in ± 2kb around TSS, the signal was log2 transformed, the differences were calculated and plotted using R.

### Normalization of AcH2AZ signal against H2AZ profile

Normalization factors were calculated for the AcH2AZ and H2AZ libraries in siControl and siTIP60 conditions with CSAW (Lun & Smyth, 2016) by counting the reads in 10kb windows along the entire genome, but excluding the blacklisted regions from ENCODE. Peaks for AcH2AZ and H2AZ libraries in siControl and siTIP60 conditions were called using Macs2 broadPeak mode with default parameters (Zhang et al, 2008). Peaks falling within the TSS +/- 2kb regions were merged and called ‘TSS peak regions’. Reads from the 4 libraries were counted on TSS peak regions using regionCounts from CSAW. Raw counts were transformed with the Rlog transformation from DESeq2 and normalized using the normalization factors previously calculated by CSAW. The AcH2AZ/H2AZ ratio was calculated for each one of the TSS peak regions in siControl and siTIP60 conditions. This process was iterated throughout the 741 TSS regions of downregulated genes after siTIP60 treatment.

### RNA-seq analysis

TIP60-depleted samples sent for RNA-seq were from 1 set each performed in MCF10A and MCF10A-MSCV. Two sets of H2AZ-depleted samples sent for RNA sequencing were performed in MCF10A. RNA-seq paired reads were aligned to the human reference genome GRCh37/hg19 and the gencode release 19 transcript database using ‘STAR Aligner’ (v2.5.1b) (Dobin et al, 2013) with default parameters. Transcript levels quantification were computed by ‘FeatureCounts’ from the ‘Rsubread’ package (v1.24) (Liao et al, 2013) with the parameter strand Specific set to 1. The transcript counts in siControl and siTIP60 experiments were subsequently subjected to differential expression analysis with ‘DESeq2’ (v1.14.1) (Love et al, 2014) using a Likelihood Ratio Test to estimate the expression changes. Differentially expressed transcripts were determined as those with an adjusted p-value lower than 0.05 and an absolute log2 fold change greater than 1. Boxplots were drawn using the read counts after applying the Variance Stabilizing Transformation. Significance of the mean difference between the distributions in the boxplots was calculated using Mann-Whitney U Test function from R. The volcano plot and heatmaps of gene expression were drawn using the scatterplot and heatmap.2 function from R/Bioconductor respectively. Functional annotation in enriched pathways of the gene list was performed with ConsensusPathDB (Kamburov et al, 2011), using the KEGG and REACTOME databases and p-adjusted values thresholds < 0.01.

### Statistical analysis

Student t-test was used for statistical comparisons unless otherwise stated in the respective figure legends. p-values are depicted as such (*, *p* < 0.05; **, *p* < 0.01; ***, *p* < 0.005). Where *p>0*.*05*, the p-value is shown. The number of independent replicates used for statistical analysis in all cases is indicated in the respective figure legends. N represents biological replicates. Error bars were plotted using either standard deviation (SD) or standard error of the mean (SEM) and are indicated in the respective figure legends.

## Data availability

The RNA-seq and ChIP-seq data from this study have been submitted to the NCBI Gene Expression Omnibus (GEO: https://www.ncbi.nlm.nih.gov/geo/query/acc.cgi?acc=GSE110407) under accession number GSE110407.

## Acknowledgments

This work was supported by grants from National Research Foundation Singapore and the Singapore Ministry of Education under its Research Centers of Excellence initiative to the Cancer Science Institute of Singapore [R-713-006-014-271 and R-713-103-006-135 to SJ]; Ministry of Education Academic Research Fund [MOE AcRF Tier 1 T1-2012 Oct -04 and T1-2016 Apr -01]; RNA Biology Center at CSI Singapore, NUS, from funding by the Singapore Ministry of Education’s Tier 3 grants [MOE2014-T3-1-006]; Post-graduate fellowship by the Cancer Science Institute of Singapore, National University of Singapore (to YZ, KKL, RTM, SSB, CYT and MMH); Post-graduate fellowship awarded by Yong Loo Lin School of Medicine, National University of Singapore (to DR).

## Author Contributions

YZ, KKL and SJ designed and conceived the experiments used in this study. YZ, KKL, DR, SSB, LN, DN, WST, CYT conducted the experiments. KKL, YZ, LN performed protein purification and HAT assay. YZ generated samples for RNA-seq and ChIP-seq, and CYT generated samples for 1 set of RNA-seq data. YZ, KKL, LN, SSB and DR validated RNA-seq results. DR and SSB validated ChIP-seq results. KKL, YZ, DR, SSB and LN performed doxorubicin sensitivity assays. KKL performed cisplatin and camptothecin sensitivity assays. DN performed knockdown in various cell lines. WST attempted some mice experiments. RTM and SSB conducted and, TB and SJ supervised the bioinformatics analysis used in this study. MMH, ADJ, YT, CWJ and DGT helped in the better design of experiments and provided valuable intellectual input.

