## SUPPLEMENTARY MATERIALS for "TIP60 acetylates H2AZ and regulates doxorubicin-induced DNA damage sensitivity through *RAD51* transcription"

### Table of contents:


- Supplementary Figure Legends
- Supplementary Figure S1
- Supplementary Figure S2
- Supplementary Figure S3
- Supplementary Figure S4
- Supplementary Figure S5
- Supplementary Figure S6
- Supplementary Figure S7
- Supplementary Figure S8
- Supplementary Figure S9
- Supplementary Figure S10
- Supplementary Figure S11
- Supplementary Figure S12
- Supplementary Figure S13
- Supplementary Figure S14
- Supplementary Figure S15
- Supplementary Figure S16
- Supplementary Figure S17
- Supplementary Figure S18

### SUPPLEMENTARY FIGURE LEGENDS

**Fig. S1 - Chromatin immunoprecipitation followed by sequencing (ChIP-seq) analysis shows TIP60 largely affects H2AZ and AcH2AZ enrichment on gene promoters.**

(**A and B**) Pie chart showing the distribution of AcH2AZ peak (A) and H2AZ peak (B) on chromatin altered by depletion of TIP60 in MCF10A. More than 40% of the peaks are localized on gene promoter. (N = 1).

**Fig. S2 - H2AZ and AcH2AZ peaks enriched specifically on the promoters of known TIP60 occupied genes.**

(**A and B**) Genome browser shot of normalized occupancy profiles for *Nucleolin* (positive target) (A) and *ACHR* (negative target) (B). For comparison between the two experiments, the scales for siControl and siTIP60 profiles are set to the same range for the respective H2AZ (purple) and AcH2AZ (blue) tracks. (N = 1).

**Fig. S3 - RNA sequencing dataset analysis of TIP60 and H2AZ depletion.**

(**A**) Volcano plot showing gene expression changes upon TIP60 depletion. 523 genes were upregulated, and 850 genes were downregulated. Genes with FDR < 0.05 and |log2 (fold change) |> 1 were considered significant change. (N = 2).

(**B**) Volcano plot showing gene expression changes upon H2AZ depletion. 525 genes were upregulated, and 436 genes were downregulated. Genes with FDR < 0.05 and |log2 (fold change) |> 1 were considered significant change. (N = 2).

**Fig. S4 - KEGG pathway enrichment of downregulated genes from RNA-seq data.**

(**A and B**) KEGG pathway enrichment analysis based only on downregulated genes from RNA-seq data analyses from TIP60-depleted (A) and H2AZ depleted (B) samples without taking into consideration changes in H2AZ or acH2AZ ChIP-seq data. The bars show the percentage of the differentially expressed gene from the total number of genes associated with a given pathway. Pathways related to cell cycle and DNA repair are highlighted in red. (N = 2).

**Fig. S5 - TIP60 regulates RAD51 expression.**

Western blot showing RAD51 protein level decrease upon TIP60 depletion using 2 different TIP60 siRNAs in MCF10A. Actin serves as a loading control. (N = 1).

**Fig. S6 - TIP60 regulates *RAD51* expression in different cell lines.**

(**A-D**) *RAD51* and *TIP60* expression upon TIP60 depletion in DLD1 (A) (N = 2), HEPG2 (B) (N = 2), HCT116 (C) (N = 3), and SNU398 (D) (N = 3). The expression was validated by qPCR and results were analyzed as fold change against control. All primers used in qPCR experiments are shown in Table 3. All data represent means ± SD. **, p-value < 0.01; ***, p-value < 0.005.

**Fig. S7 - Validation of genes enriched in cell cycle and DNA repair pathways.**

(**A and B**) Expression of genes involved in homologous recombination and cell cycle upon TIP60 (A) or H2AZ (B) depletion in MCF10A. The expression was validated by qPCR and results were analyzed as fold change against control. All primers used in qPCR experiments are shown in Table 3. NC, Non-targeting Control. (N = 3). All data represent means ± SD. n.s., p > 0.05; *, p < 0.05; **, p < 0.01; ***, p < 0.005.

**Fig. S8 - TIP60 regulates AcH2AZ/H2B level on *RAD51* promoter.**

(**A and B**) ChIP-qPCR showing occupancy of H2B (A) and AcH2AZ/H2B ratio (B) on *RAD51* TSS upon TIP60 depletion in MCF10A (N = 3). All data represent means ± SEM.

**Fig. S9 - TIP60 regulates mainly AcH2AZ occupancy on *ANP32E*, *ESPL1*, and *E2F2* promoters.**

(**A-E**) ChIP-qPCR validation showing occupancy of AcH2AZ/H2AZ ratio (A), AcH2AZ/H2B ratio (B), AcH2AZ (C), H2AZ (D), and H2B (E) on *ANP32E*, *ESPL1*, *E2F2*, and *ACHR* TSS upon TIP60 depletion in MCF10A. (N = 3). All data represent means ± SEM. *, p-value < 0.05; **, p-value < 0.01.

**Fig. S10 - TIP60 enriches on *RAD51* promoter**.

(**A-C**) IGV screenshots of TIP60 localization on the *RAD51* promoter from publicly available ChIP-seq datasets of mouse embryonic stem cells GSE69671 (A) and GSM1650007 (B), and FLAG-TIP60 localization on the *RAD51* promoter from K562 cell line GSE69645 (C). TIP60 localization peaks on the *RAD51* promoter are marked with a red square.

(**D**) FLAG-TIP60 ChIP-qPCR showing occupancy of TIP60 on *RAD51* promoter in MCF10A stably expressing FLAG-TIP60. All data represent means ± SD. SD represents standard deviation from technical replicates. Specificity of the signal is shown with TIP60 depletion. (N = 1).

**Fig. S11 - TIP60-dependent doxorubicin sensitivity is dependent on RAD51 level**.

(**A and B**) Doxorubicin sensitivity assay showing rescue in cell viability in RAD51 expressing MCF10A cells depleted of TIP60 and treated with indicated amounts of doxorubicin from biological sets 2 (A) and 3 (B). Cell viability was measured using an MTS assay. All data represent means ± SD. SD represents standard deviation from technical replicates. p-value was calculated using technical replicates within the set. **, p-value < 0.01; ***, p < 0.005.

**Fig. S12 - H2AZ-dependent doxorubicin sensitivity is dependent on RAD51 level.**

(**A and B**) Doxorubicin sensitivity assay showing rescue in cell viability in RAD51 expressing MCF10A cells depleted of H2AZ and treated with indicated amounts of doxorubicin from biological sets 1 (A) and 2 (B). Cell viability was measured using an MTS assay. All data represent means ± SD. SD represents standard deviation from technical replicates. p-value was calculated using technical replicates within the set. **, p-value < 0.01; ***, p < 0.005.

**Fig. S13 – TIP60-dependent camptothecin sensitivity is dependent on RAD51 level.**

Camptothecin sensitivity assay showing rescue in cell viability in RAD51 expressing MCF10A cells depleted of TIP60 and treated with indicated amounts of camptothecin. (N = 2). Cell viability was measured using an MTS assay. All data represent means ± SEM. *, p-value < 0.05.

**Fig. S14 - TIP60/H2AZ-dependent sensitivity to cisplatin is not dependent on RAD51 level.**

(**A and B**) Cisplatin sensitivity assay shows no rescue in cell viability in RAD51 expressing MCF10A cells depleted of TIP60 (A) (N = 3) or H2AZ (B) (N = 2) and treated with indicated amounts of cisplatin. Cell viability was measured using an MTS assay. All data represent means ± SEM.

**Fig. S15 - Majority of genes regulated by TIP60 overlaps with AcH2AZ changes.**

Overlap strategy without including H2AZ depletion RNA-seq dataset shows 741 upregulated genes and 361 downregulated genes regulated by TIP60-dependent H2AZ acetylation. Dots marked in red represent genes from the overlap datasets.

**Fig. S16 - H2AZ regulated DNA repair and cell cycle gene expression is mainly through H2AZ.2.**

Expression of genes involved in homologous recombination and cell cycle upon H2AZ.1 alone, H2AZ.2 alone, or H2AZ.1 and H2AZ.2 depletion in MCF10A. The expression was validated by qPCR and results were analyzed as fold change against control. All primers used in qPCR experiments are shown in Table 3. (N = 3). All data represent means ± SD. NS, $p > 0.05$; *, $p < 0.05$; **, $p < 0.01$; ***, $p < 0.005$.

**Fig. S17 - E2F1 and E2F4 are predicted to bind to *RAD51* promoter and regulate *RAD51* expression.**

List of transcription factors predicted to bind to promoter sites of the 66 genes regulated by TIP60 and H2AZ and which regulates *RAD51* utilizing the Transcriptional Regulatory Relationships Unraveled by Sentence-based Text mining (TRRUST) tool.

**Fig. S18 - TIP60 may regulate H2AZ transcriptionally.**

(**A and B**) Relative expression of *H2AZ.1* and *H2AZ.2* levels upon TIP60 depletion in MCF10A from biological set 1 (A) and set 2 (B). All data represent means ± SD. SD represents standard deviation from technical replicates.

(**C and D**) IGV screenshots showing AcH2AZ and H2AZ occupancy upon TIP60 depletion on *H2AFZ* (C) and *H2AFV* (D) TSS. AcH2AZ and H2AZ peaks on the TSS of *H2AFZ* and *H2AFV* are marked with a red box.

**Fig. S1**

**A**

*siTIP60* vs *siControl*

Distribution of AcH2AZ enrichment on chromatin

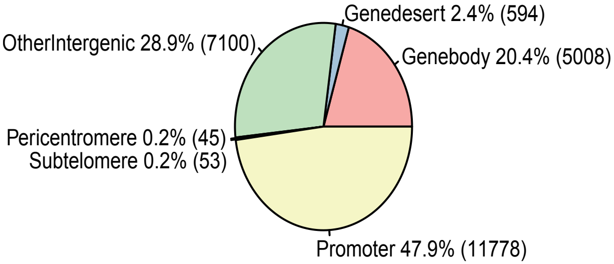


**B**

*siTIP60* vs *siControl*

Distribution of H2AZ enrichment on chromatin

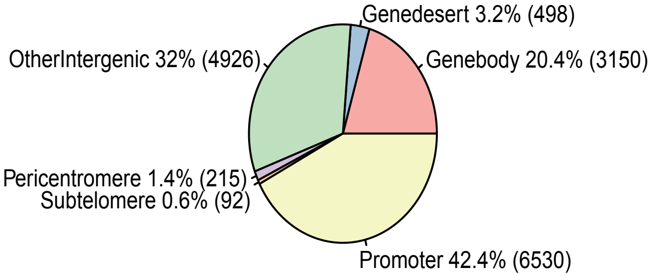

Fig. S2

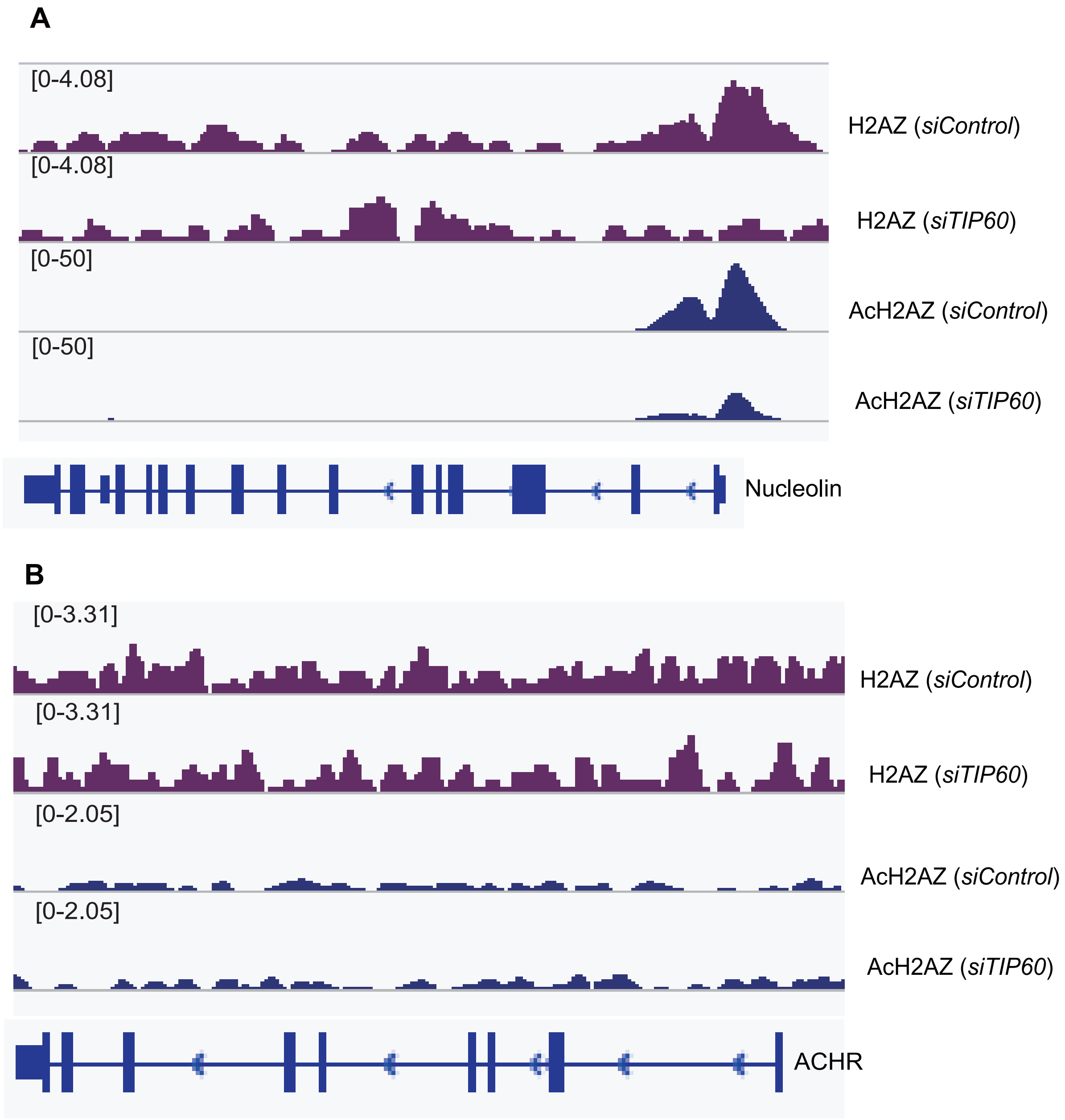
A
[0-4.08]
H2AZ (siControl)
[0-4.08]
H2AZ (siTIP60)
[0-50]
AcH2AZ (siControl)
[0-50]
AcH2AZ (siTIP60)
Nucleolin
B
[0-3.31]
H2AZ (siControl)
[0-3.31]
H2AZ (siTIP60)
[0-2.05]
AcH2AZ (siControl)
[0-2.05]
AcH2AZ (siTIP60)
ACHR

Fig. S3

A

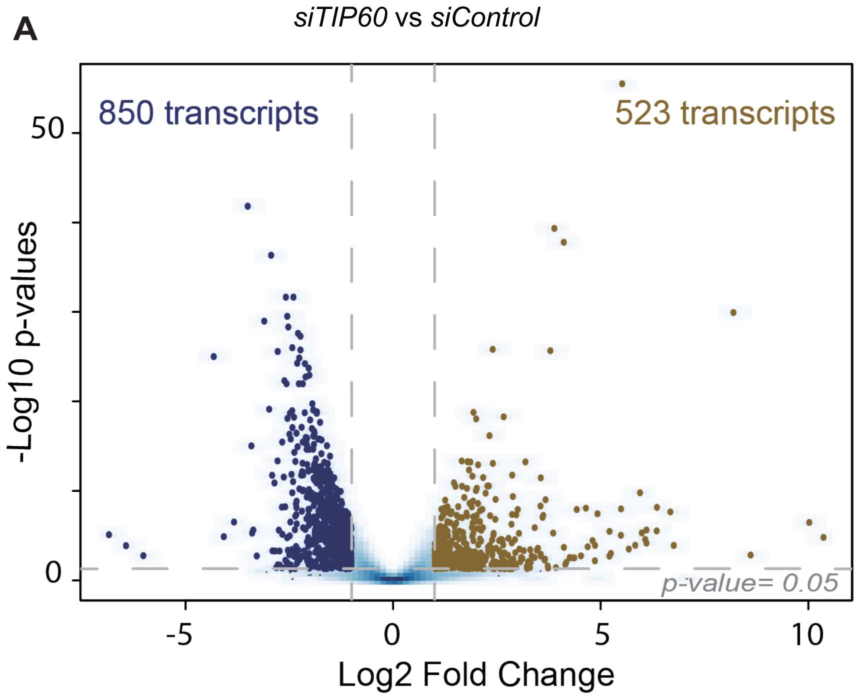
siTIP60 vs siControl
850 transcripts
523 transcripts
50
0
-Log10 p-values
p-value= 0.05
-5
0
5
10
Log2 Fold Change

B

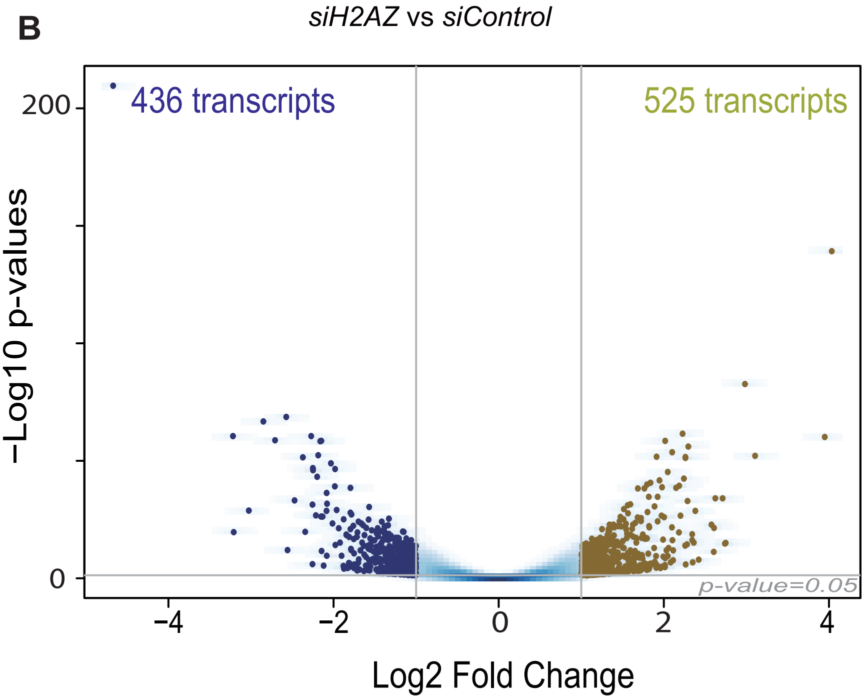
siH2AZ vs siControl
436 transcripts
525 transcripts
200
0
−Log10 p-values
p-value=0.05
−4
−2
0
2
4
Log2 Fold Change

Fig. S4

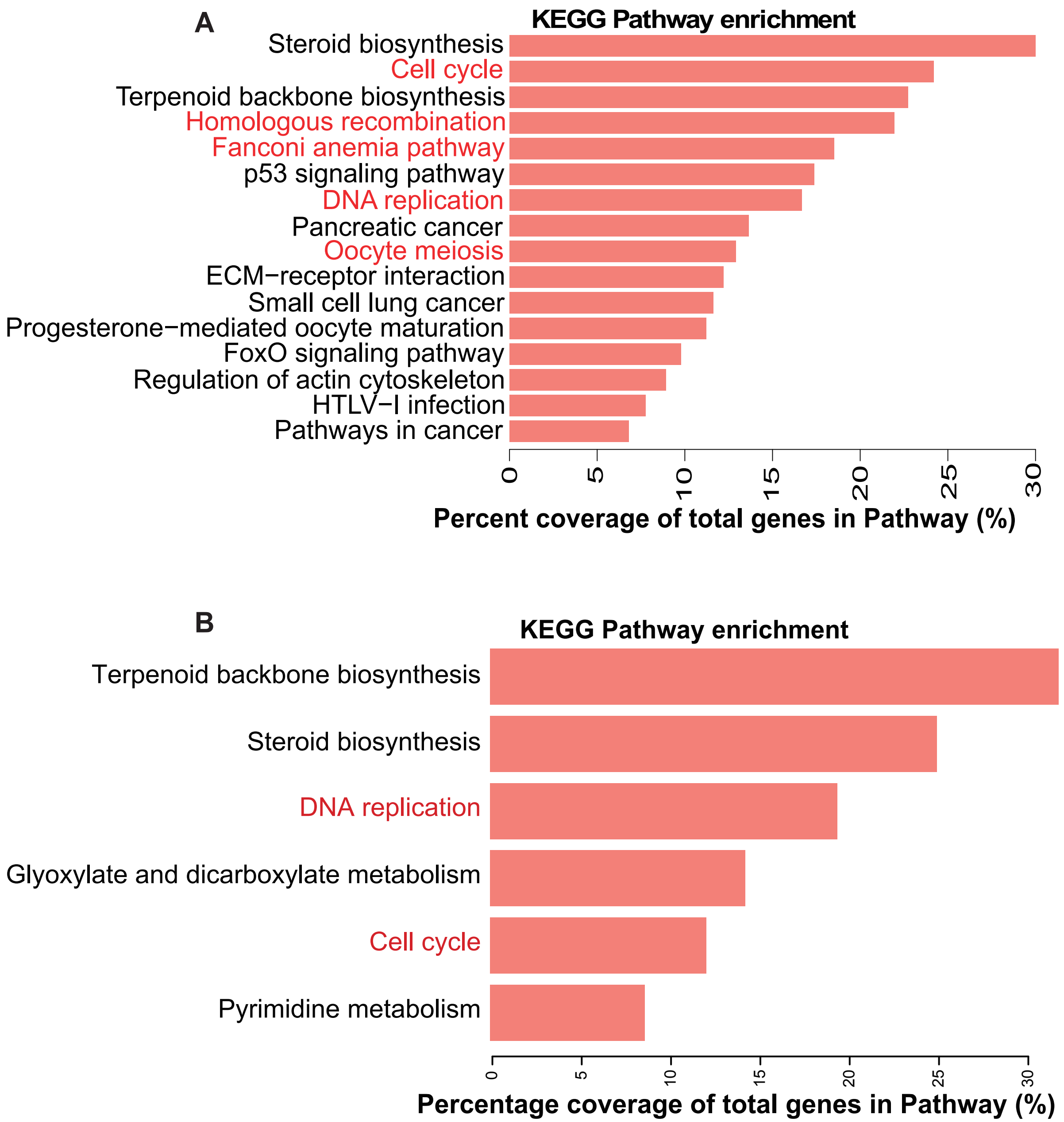
A
KEGG Pathway enrichment
Steroid biosynthesis
Cell cycle
Terpenoid backbone biosynthesis
Homologous recombination
Fanconi anemia pathway
p53 signaling pathway
DNA replication
Pancreatic cancer
Oocyte meiosis
ECM−receptor interaction
Small cell lung cancer
Progesterone−mediated oocyte maturation
FoxO signaling pathway
Regulation of actin cytoskeleton
HTLV−I infection
Pathways in cancer
0
5
10
15
20
25
30
Percent coverage of total genes in Pathway (%)
B
KEGG Pathway enrichment
Terpenoid backbone biosynthesis
Steroid biosynthesis
DNA replication
Glyoxylate and dicarboxylate metabolism
Cell cycle
Pyrimidine metabolism
0
5
10
15
20
25
30
Percentage coverage of total genes in Pathway (%)

Fig. S5

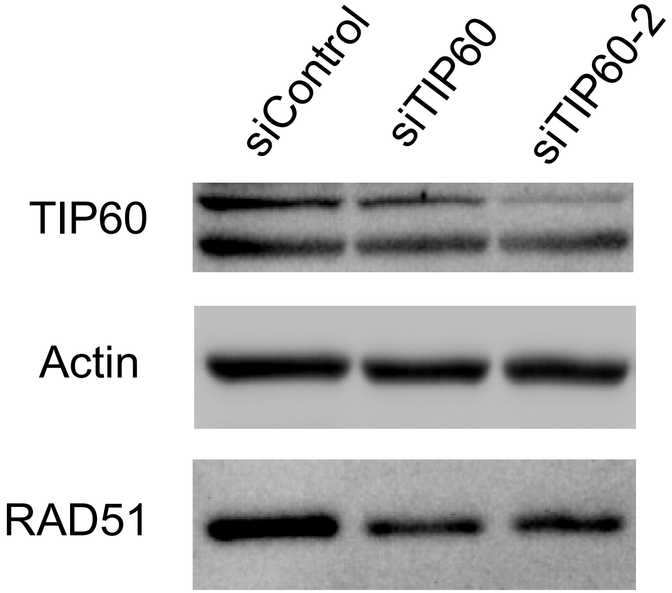

**Fig. S6**

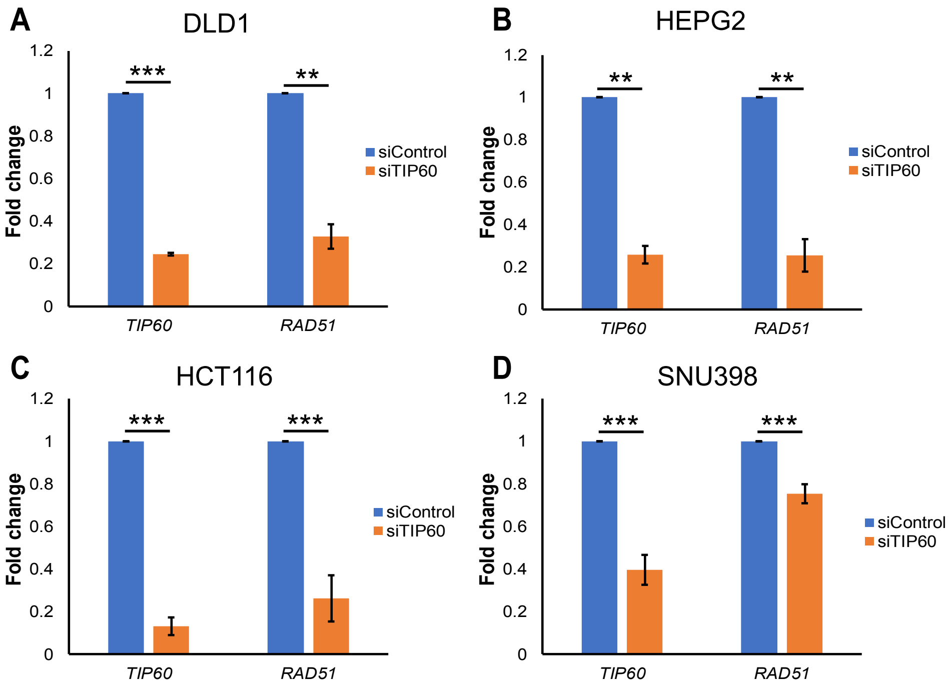
A
DLD1
B
HEPG2
C
HCT116
D
SNU398
Fold change
1.2
1
0.8
0.6
0.4
0.2
0
***
**
siControl
siTIP60
TIP60
RAD51

Fig. S7

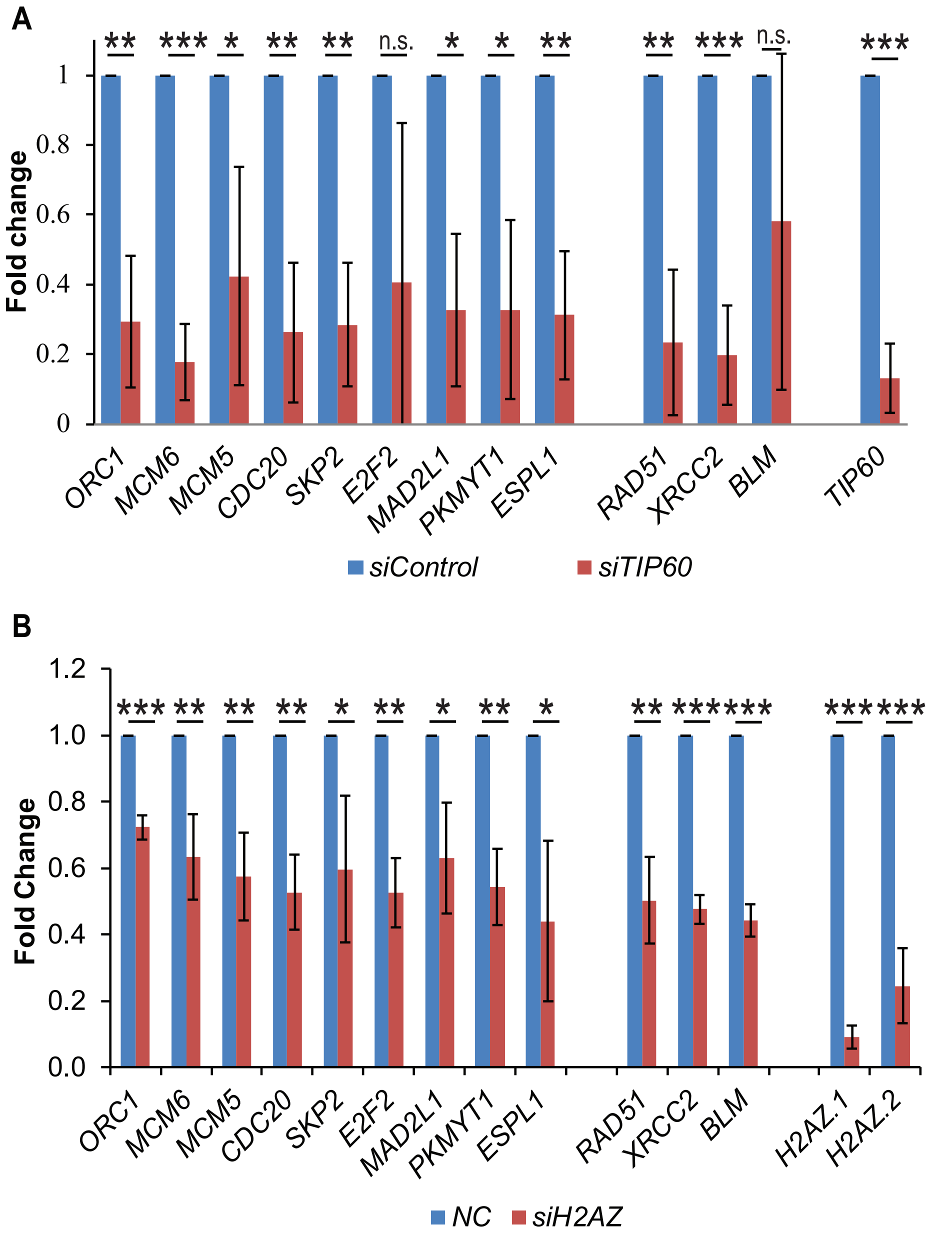
A
Fold change
1
0.8
0.6
0.4
0.2
0
**
***
*
**
**
n.s.
*
*
**
**
***
n.s.
***
ORC1
MCM6
MCM5
CDC20
SKP2
E2F2
MAD2L1
PKMYT1
ESPL1
RAD51
XRCC2
BLM
TIP60
siControl
siTIP60
B
Fold Change
1.2
1.0
0.8
0.6
0.4
0.2
0.0
***
**
**
**
*
**
*
**
*
**
***
***
***
***
ORC1
MCM6
MCM5
CDC20
SKP2
E2F2
MAD2L1
PKMYT1
ESPL1
RAD51
XRCC2
BLM
H2AZ.1
H2AZ.2
NC
siH2AZ

Fig. S8

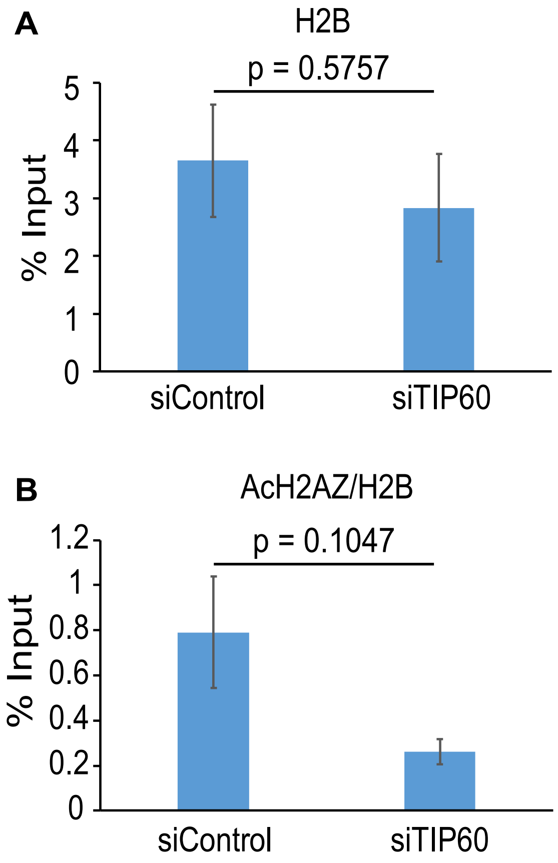
A
H2B
p = 0.5757
% Input
5
4
3
2
1
0
siControl
siTIP60
B
AcH2AZ/H2B
p = 0.1047
% Input
1.2
1
0.8
0.6
0.4
0.2
0
siControl
siTIP60

Fig. S9

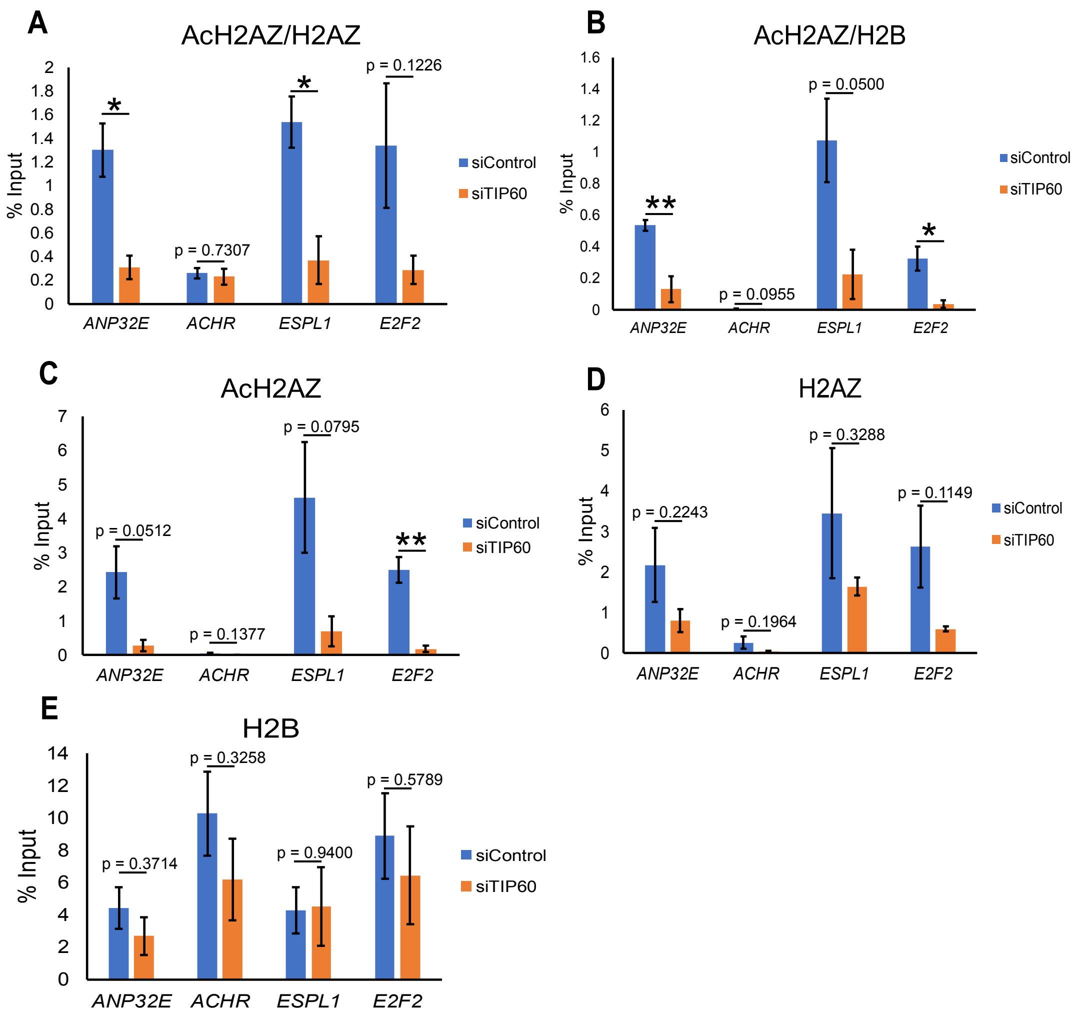

A
AcH2AZ/H2AZ
% Input
2
1.8
1.6
1.4
1.2
1
0.8
0.6
0.4
0.2
0
*
p = 0.7307
*
p = 0.1226
ANP32E
ACHR
ESPL1
E2F2
siControl
siTIP60
B
AcH2AZ/H2B
% Input
1.6
1.4
1.2
1
0.8
0.6
0.4
0.2
0
**
p = 0.0955
p = 0.0500
*
ANP32E
ACHR
ESPL1
E2F2
siControl
siTIP60
C
AcH2AZ
% Input
7
6
5
4
3
2
1
0
p = 0.0512
p = 0.1377
p = 0.0795
**
ANP32E
ACHR
ESPL1
E2F2
siControl
siTIP60
D
H2AZ
% Input
6
5
4
3
2
1
0
p = 0.2243
p = 0.1964
p = 0.3288
p = 0.1149
ANP32E
ACHR
ESPL1
E2F2
siControl
siTIP60
E
H2B
% Input
14
12
10
8
6
4
2
0
p = 0.3714
p = 0.3258
p = 0.9400
p = 0.5789
ANP32E
ACHR
ESPL1
E2F2
siControl
siTIP60

Fig. S10

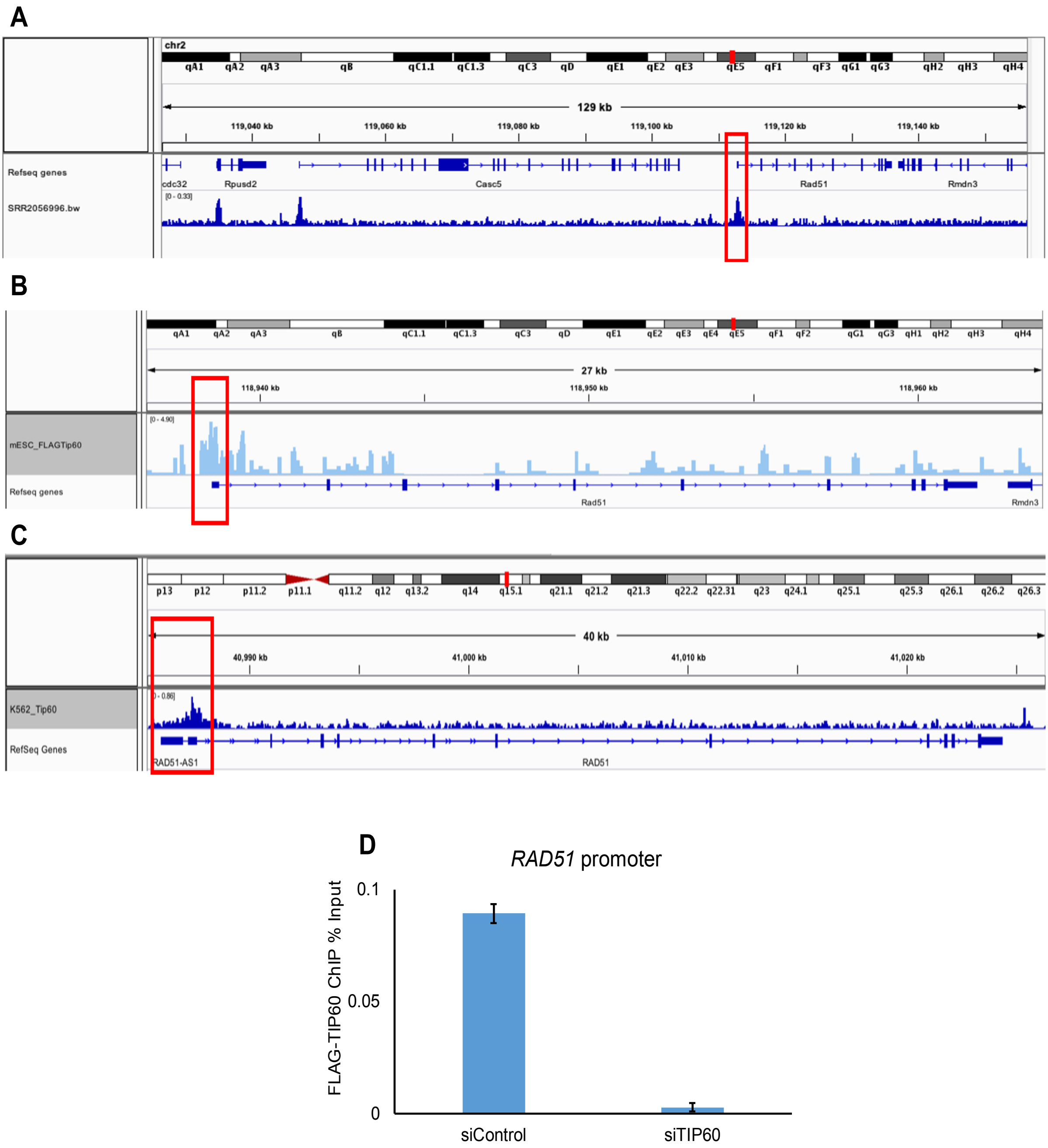
A
chr2
129 kb
119,040 kb
119,060 kb
119,080 kb
119,100 kb
119,120 kb
119,140 kb
Refseq genes
SRR2056996.bw
cdc32
Rpusd2
Casc5
Rad51
Rmdn3
B
27 kb
118,940 kb
118,950 kb
118,960 kb
mESC_FLAGTip60
Refseq genes
Rad51
Rmdn3
C
40 kb
40,990 kb
41,000 kb
41,010 kb
41,020 kb
K562_Tip60
RefSeq Genes
RAD51-AS1
RAD51
D
RAD51 promoter
FLAG-TIP60 ChIP % Input
0.1
0.05
0
siControl
siTIP60

Fig. S11

A

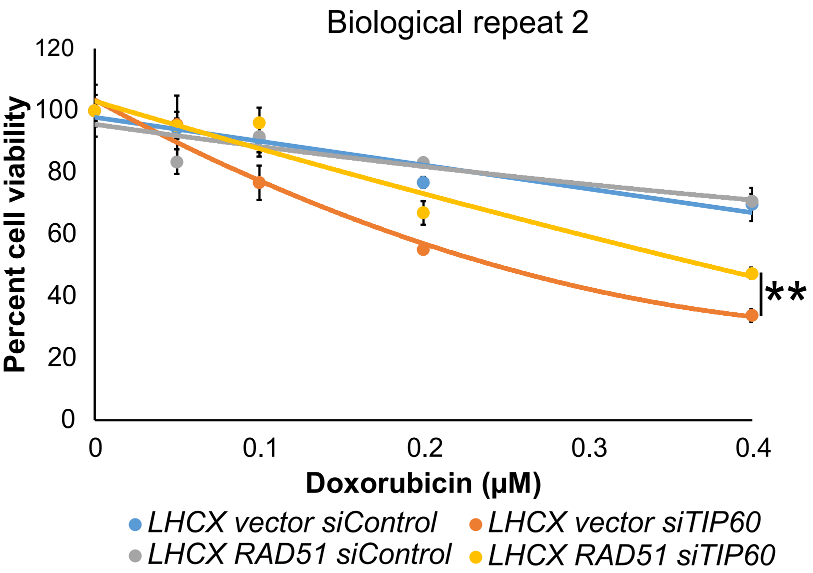

Biological repeat 2
120
100
80
60
40
20
0
Percent cell viability
0
0.1
0.2
0.3
0.4
Doxorubicin (µM)
**
LHCX vector siControl
LHCX vector siTIP60
LHCX RAD51 siControl
LHCX RAD51 siTIP60


B

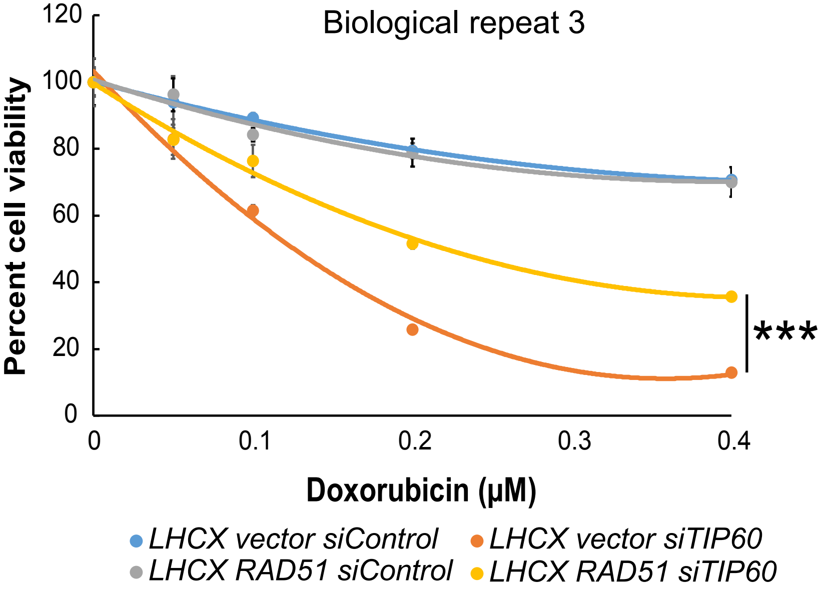

Biological repeat 3
120
100
80
60
40
20
0
Percent cell viability
0
0.1
0.2
0.3
0.4
Doxorubicin (µM)
***
LHCX vector siControl
LHCX vector siTIP60
LHCX RAD51 siControl
LHCX RAD51 siTIP60

Fig. S12

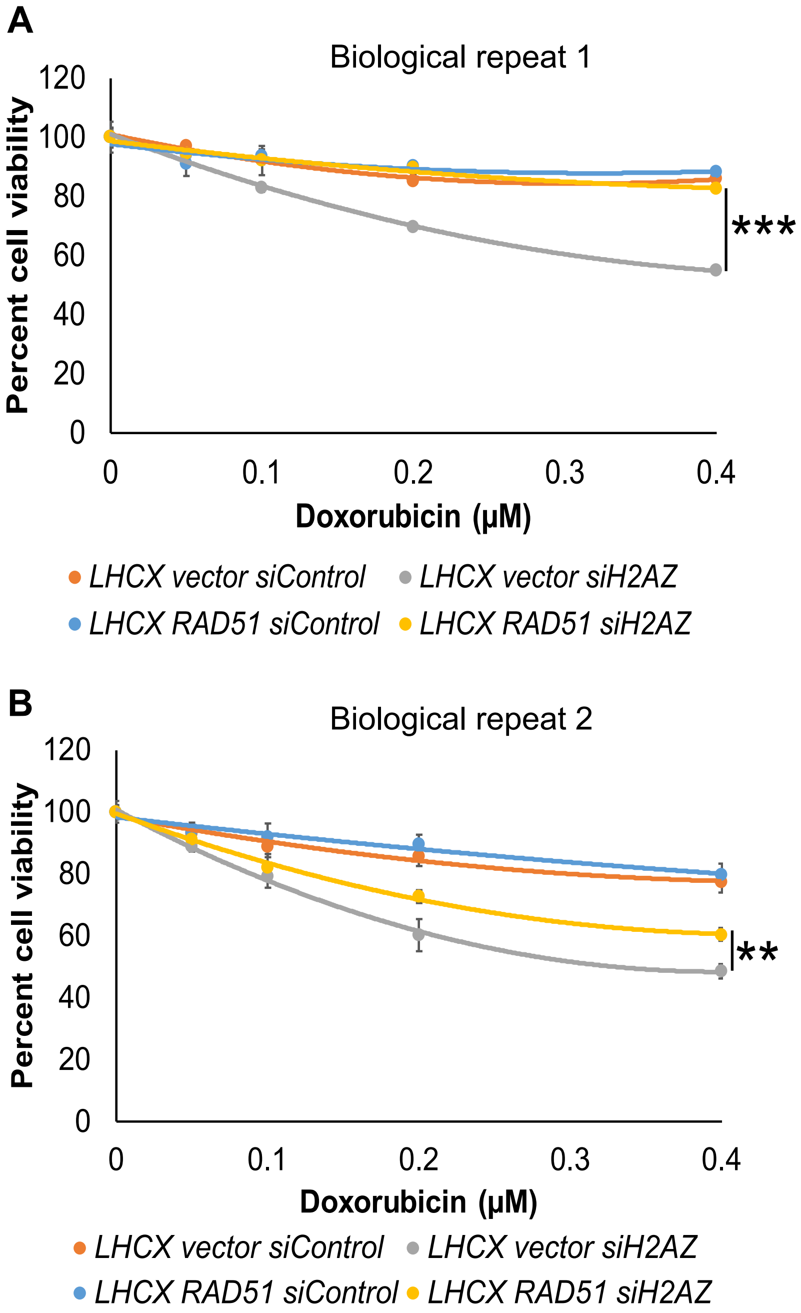
A
Biological repeat 1
Percent cell viability
120
100
80
60
40
20
0
0
0.1
0.2
0.3
0.4
Doxorubicin (μM)
***
LHCX vector siControl
LHCX vector siH2AZ
LHCX RAD51 siControl
LHCX RAD51 siH2AZ
B
Biological repeat 2
Percent cell viability
120
100
80
60
40
20
0
0
0.1
0.2
0.3
0.4
Doxorubicin (μM)
**
LHCX vector siControl
LHCX vector siH2AZ
LHCX RAD51 siControl
LHCX RAD51 siH2AZ

**Fig. S13**

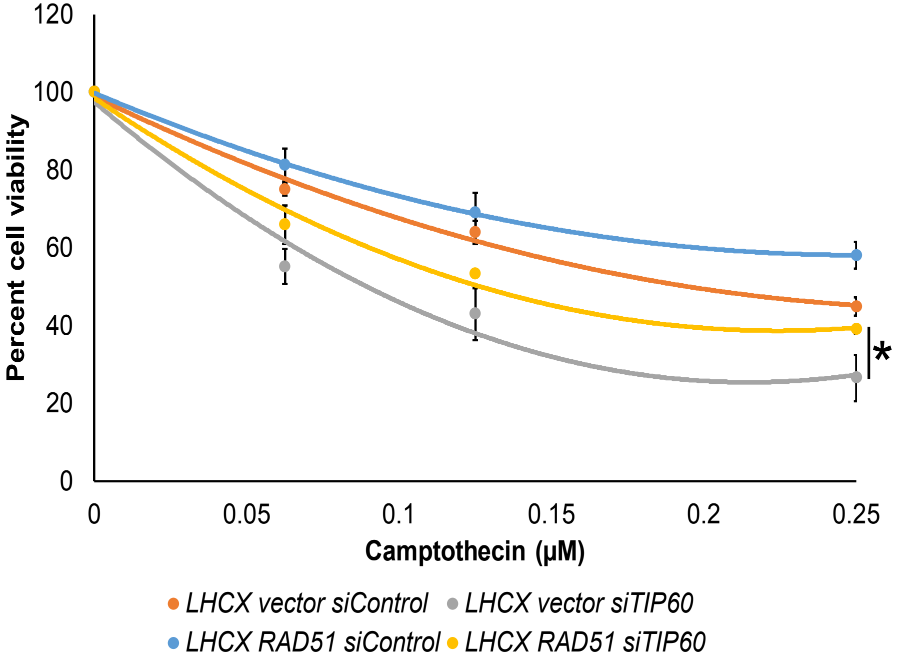
120
100
80
60
40
20
0
Percent cell viability
0
0.05
0.1
0.15
0.2
0.25
Camptothecin (μM)
*
LHCX vector siControl
LHCX vector siTIP60
LHCX RAD51 siControl
LHCX RAD51 siTIP60

**Fig. S14**

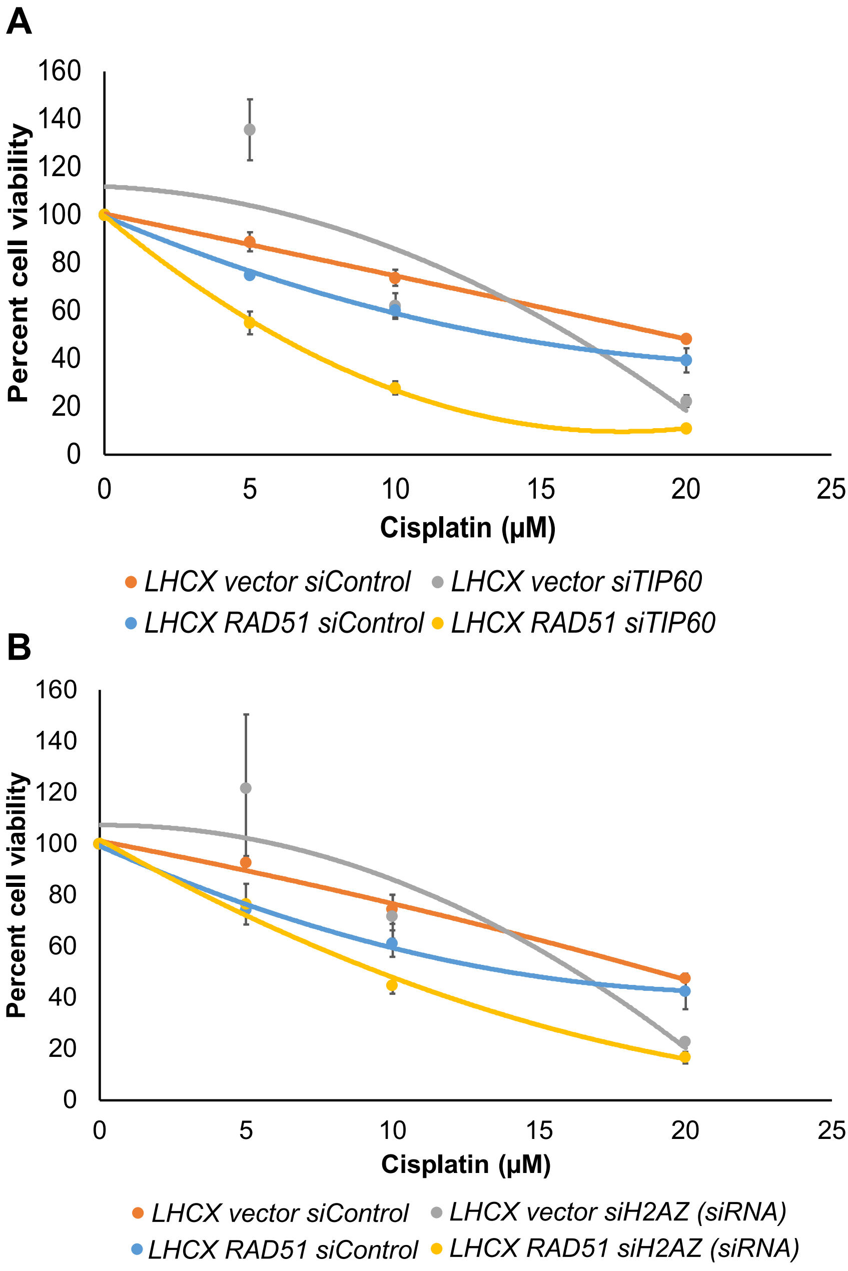
A
Percent cell viability
160
140
120
100
80
60
40
20
0
0
5
10
15
20
25
Cisplatin (µM)
LHCX vector siControl
LHCX vector siTIP60
LHCX RAD51 siControl
LHCX RAD51 siTIP60
B
Percent cell viability
160
140
120
100
80
60
40
20
0
0
5
10
15
20
25
Cisplatin (µM)
LHCX vector siControl
LHCX vector siH2AZ (siRNA)
LHCX RAD51 siControl
LHCX RAD51 siH2AZ (siRNA)

**Fig. S15**

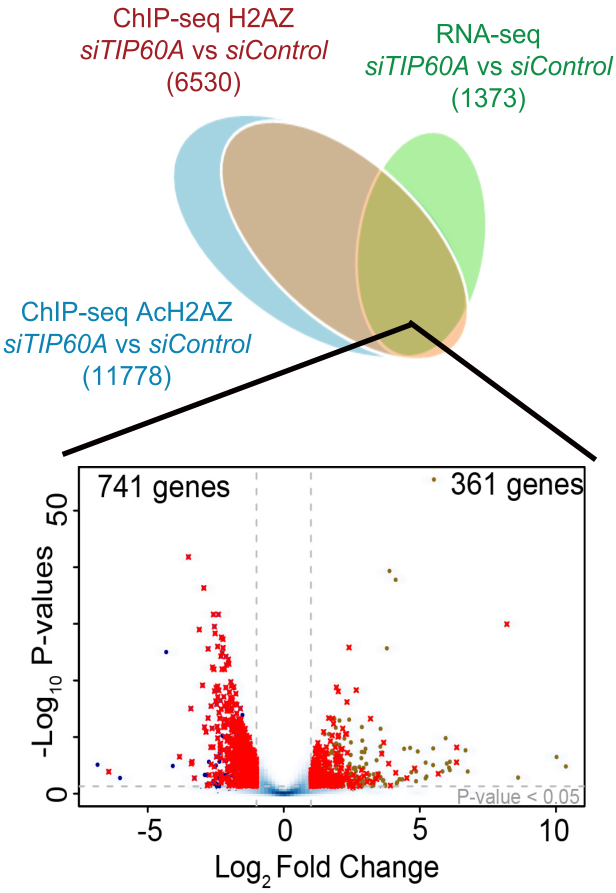
ChIP-seq H2AZ
*siTIP60A* vs *siControl*
(6530)
RNA-seq
*siTIP60A* vs *siControl*
(1373)
ChIP-seq AcH2AZ
*siTIP60A* vs *siControl*
(11778)
741 genes
361 genes
$-\log_{10}$ P-values
P-value < 0.05
0
50
-5
0
5
10
$\log_2$ Fold Change

**Fig. S16**

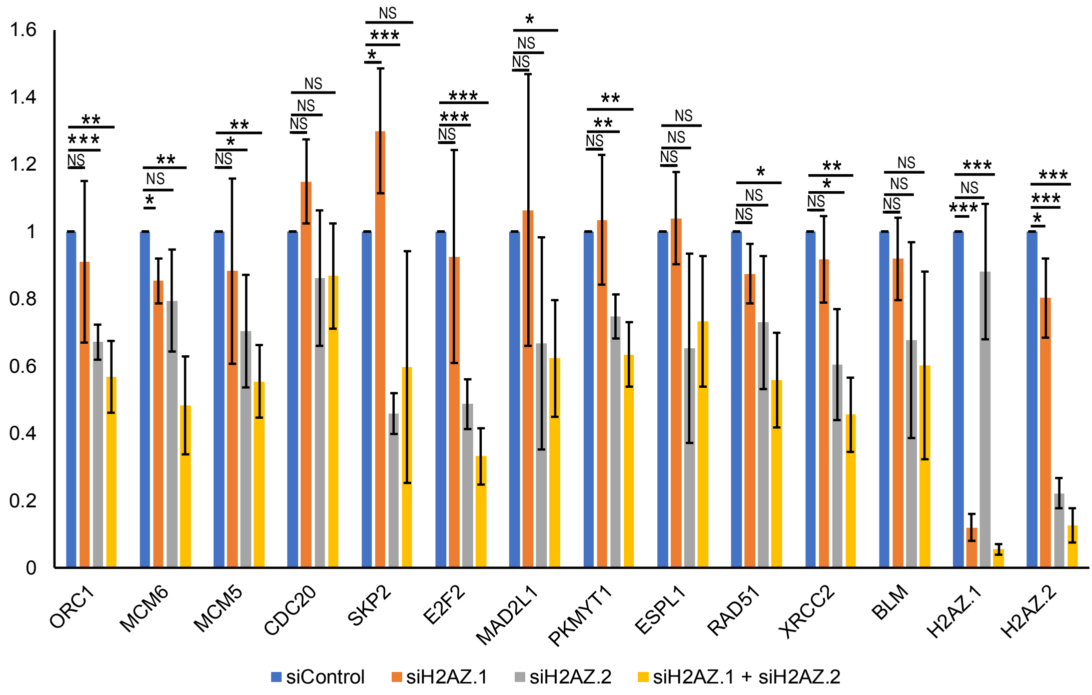

1.6
1.4
1.2
1
0.8
0.6
0.4
0.2
0
ORC1
MCM6
MCM5
CDC20
SKP2
E2F2
MAD2L1
PKMYT1
ESPL1
RAD51
XRCC2
BLM
H2AZ.1
H2AZ.2
siControl
siH2AZ.1
siH2AZ.2
siH2AZ.1 + siH2AZ.2
NS
*
**
***

Fig. S17

| Key TF | Description | List of overlapped genes |
| --- | --- | --- |
| E2F1 | E2F transcription factor 1 | MYBL2,MCM5,THBS1,SKP2,RRM2, RRP1B,TYMS,RAD51,UHRF1 |
| E2F4 | E2F transcription factor 4, p107/p130-binding | MCM10,RAD51 |

**Figure S18**

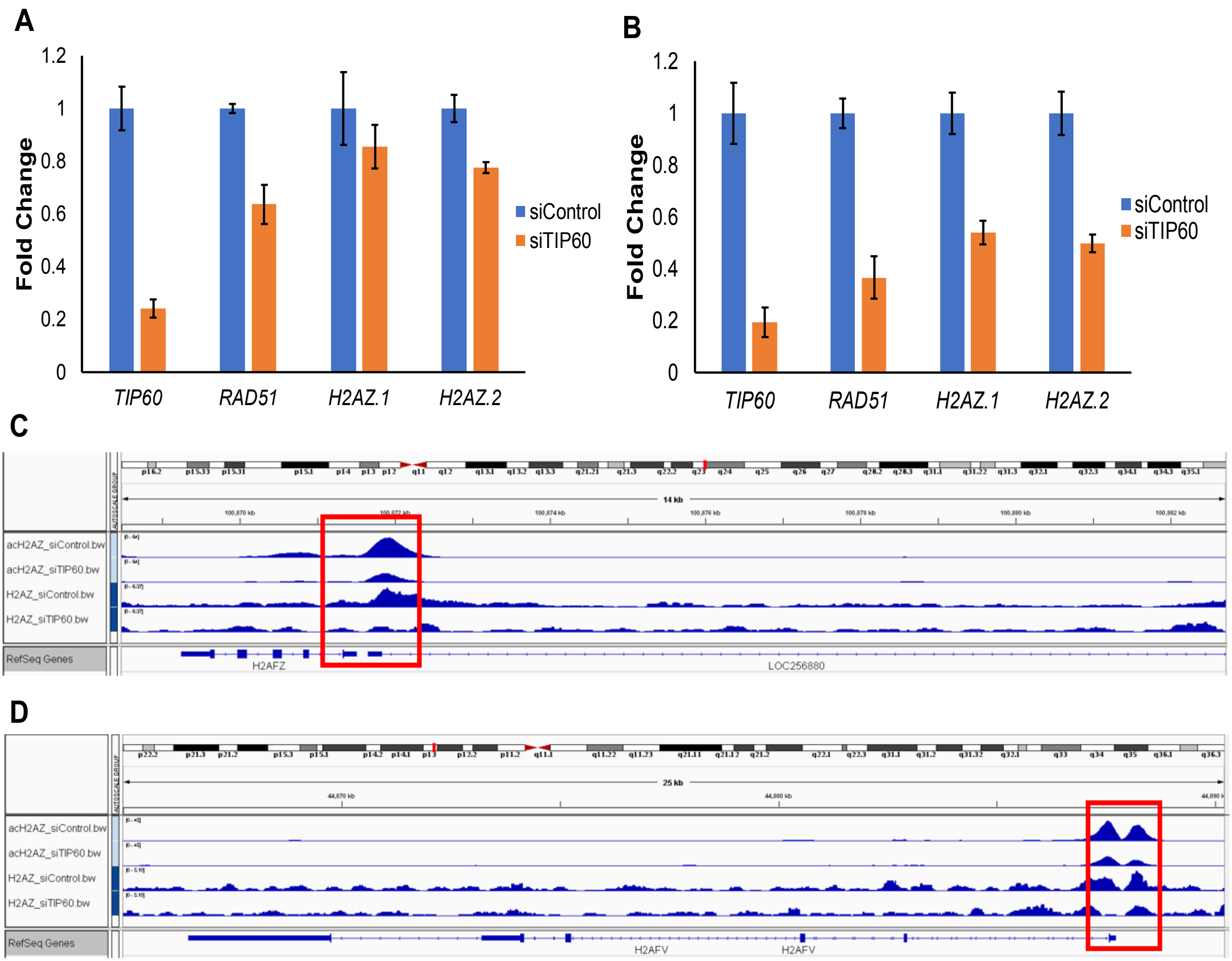
A
Fold Change
1.2
1
0.8
0.6
0.4
0.2
0
siControl
siTIP60
TIP60
RAD51
H2AZ.1
H2AZ.2
B
Fold Change
1.2
1
0.8
0.6
0.4
0.2
0
siControl
siTIP60
TIP60
RAD51
H2AZ.1
H2AZ.2
C
14 kb
acH2AZ_siControl.bw
acH2AZ_siTIP60.bw
H2AZ_siControl.bw
H2AZ_siTIP60.bw
RefSeq Genes
H2AFZ
LOC256880
D
25 kb
acH2AZ_siControl.bw
acH2AZ_siTIP60.bw
H2AZ_siControl.bw
H2AZ_siTIP60.bw
RefSeq Genes
H2AFV
H2AFV
